# Harnessing Dye-induced Photothermal Confinement in Lipid Membranes: A Path to NIR-modulated Artificial Synaptic Vesicles

**DOI:** 10.1101/2024.07.20.604262

**Authors:** Satya Ranjan Sarker, Takeru Yamazaki, Keitaro Sou, Ichiro Takemura, Yusuke Kurita, Kayoko Nomura, Mari Ichimura, Takahito Suzuki, Ayumi Kai, Takumi Araki, Shinnosuke Hattori, Taniyuki Furuyama, Young-Tae Chang, Taketoshi Kiya, Satoshi Arai

**Affiliations:** WPI Nano Life Science Institute, Kanazawa University, Kakuma-machi, Kanazawa 920-1192, Japan; Waseda Research Institute for Science and Engineering, Waseda University, Tokyo 169-8555, Japan; Sony Semiconductor Solutions Corporation, Kanagawa 243-0014, Japan; NanoMaterials Research Institute, Kanazawa University, Kakuma-machi, Kanazawa 920-1192, Japan; Department of Chemistry, Pohang University of Science and Technology (POSTECH), Pohang, Gyeongbuk, 37673, South Korea; Graduate School of Natural Science & Technology, Kanazawa University, Kakuma-machi, Kanazawa 920-1192, Japan

**Author notes:** **Correspondence:** Satoshi Arai. These authors contributed equally to this work (Satya Ranjan Sarker, Takeru Yamazaki, Keitaro Sou, Ichiro Takemura). cellmoxa inc. Tokyo 160-0022, Japan.

## Abstract

Optical heating coupled with near-infrared (NIR) light and photothermal materials enables thermal confinement within biospecimens, minimizing undesirable thermal damage. Here, we demonstrated that photothermally heating lipid bilayers embedded with a unique phthalocyanine dye (VPc) efficiently perturbs the bilayers, resulting in increased permeability. Notably, microscopic studies revealed that the mechanism causing changes in membrane permeability may not follow the conventional temperature-sensitive liposome model. Furthermore, the heat generated by NIR laser illumination rarely diffused into the surrounding environment, and the dye was located within the bilayers at the molecular level, where it effectively transferred heat to the lipid bilayer. We prepared VPc-embedded liposomes encapsulating acetylcholine (ACh) and demonstrated the NIR laser-triggered release of ACh, creating a concentration jump across a few cells or within a limited single cell region. This method induced Ca^2+^ flux through ACh receptor stimulation in thermally delicate biospecimens such as C2C12 myotubes and the *Drosophila* brain.

## Introduction

Thermal management, including quantitative heating at a microscopic scale, has attracted attention in the fields of bioengineering and life sciences (1). Controlled heating technology, termed local hyperthermia, has provided an effective therapy to combat cancers owing to the thermal vulnerability characteristics of cancer cells (2, 3). In recent years, beyond the deactivation of cellular functions, this technology has been extended to the thermal manipulation of neuronal functions in the brain and muscles as a nongenetic modulation method (4–6). Furthermore, the activation of thermogenesis through the local heating of beige fat has been proposed as a potential therapy for obesity (7). Another approach is the use of temperature variation as an external physical stimulus to operate an effective drug delivery system (DDS) (8–10). This approach could enable the rapid release of drugs to targeted locations, circumventing slow drug release in conventional DDS. For these applications, the optical heating method offers superior spatial and temporal resolutions for precise remote temperature control (11, 12).

The principle underlying optical heating methods relies on the photothermal effect, wherein light energy is converted into heat (13). Photothermal conversion is classified into distinct pathways, including surface plasmon resonance in specific metallic nanomaterials (14, 15), nonradiative relaxation in semiconductors (16, 17), and thermal vibration in organic molecules (18, 19). Metallic and semiconducting nanomaterials exhibit excellent photothermal conversion efficiency with high photostability. Targeted heating of biospecimens, referred to as “thermal confinement,” holds great significance in expanding potential biological applications (2, 20, 21). Metallic and semiconducting nanomaterials achieve highly localized heating of target specimens, minimizing temperature increases in the surrounding environment, such as in cell culture medium (22). Despite their promising applicability, these approaches face difficulties in precisely positioning photothermal nanomaterials in proximity to the target location. In addition, controlling their placement at the molecular level can be challenging. Photothermal organic dyes have distinct advantages over other methods owing to their molecular size, which allows precise targeting. However, the heat generated by a single photothermal dye (PTD) may not always provide efficient physical perturbation of the target unless the dye accumulates densely in the particle formulation (23). Thus, the precise location of the dye can be inferred to be crucial for facilitating efficient heat transfer to target biomolecules.

This study aimed to establish thermal confinement in a lipid membrane using a PTD and subsequently alter the dynamics of the bilayer. To date, several photothermally triggerable liposomes have been documented, wherein photothermal materials, such as organic dyes or gold nanoparticles, are either encapsulated within the core or embedded into bilayers (24). In both cases, two possible mechanisms have been proposed to induce payload leakage: photothermal materials heating the surrounding media in the bulk state upon illumination, leading to membrane rupture or phase transition, and local heating to directly perturb the lipid bilayer (25). The latter has garnered attention as an effective local heating method. However, despite successful animal studies employing light-triggerable liposomes, the precise events occurring within the membranes during photothermal heating at the microscopic scale remain unclear. While a temperature increase in the media is hypothesized not to accompany ultimate specific heating in the membrane, direct observations of such phenomena are scarce, and demonstrations of effective applications highlighting this feature are even fewer (26).

To explore this concept, we examined the potential for efficient heat transfer from a PTD to establish thermal confinement within the lipid membrane while maintaining a negligible temperature rise. Subsequently, we extended this study to demonstrate its applicability as an effective delivery system, facilitating the rapid release of bioactive compounds, such as neurotransmitters, with minimal thermal damage to the biospecimens. Despite being a well-documented topic, we explored the distinct characteristics that have been overlooked in numerous studies on thermally sensitive liposome systems.

## Results

### Evaluation of photothermal dye-embedded liposomes in bulk solution

Our study intended to explore the potential of inducing phase transitions in lipid bilayers under bulk conditions through photothermal heating using a dye. To achieve this, we prepared liposomes incorporating a PTD into the lipid bilayers (Fig. 1A,B). The lipid formulation comprised 1,2-dipalmitoyl-sn-glycero-3-phosphocholine (DPPC), L-glutamic acid, *N*-(3-carboxy-1-oxopropyl)-,1,5-dihexadecyl ester (SA), and 1,2-distearoyl-sn-glycero-3-phosphoethanolamine-(polyethylene glycol) (DSPE-PEG) at a molar ratio of 9:1:0.06. According to our previous study, the phase transition temperature (*T*_c_) from gel to liquid crystalline in this mixed lipid closely resembled that of pure DPPC, which is approximately 41 °C (27). While the negatively charged lipid (SA) had little impact on the *T*_c_ of DPPC, it was expected to reduce the lamellarity of the bilayers due to electronic repulsion between the layers (27). Subsequently, we selected PTDs absorbing near-infrared (NIR) light of approximately 800 nm, including phthalocyanine vanadium oxide complex (VPc), naphthalocyanine vanadium oxide complex (VNPc), IR-792, indocyanine green (ICG), 1,1′-diethyl-2-2′-dicarbo-cyanine (DDI), and C6TI (Fig. 1A). These dyes were well-suited for further experiments using a common NIR laser operating at 808 nm (Fig. S1). Metallophthalocyanines and naphthalocyanines are well-recognized photosensitizers used in photothermal therapy (19). Specifically, vanadium oxide complexes such as VPc and VNPc are considered suitable because of their high photothermal efficiency and reduced generation of reactive oxygen species (28, 29). Additionally, linear cyanine dyes and their derivatives, such as ICG, IR792, and DDI, are frequently used to develop photothermally triggerable lipid nanoparticles (30). In addition to these dyes, a unique NIR dye, C6TI, was synthesized according to a previous study (31). To our knowledge, C6TI exhibits the highest photothermal conversion efficiency (c.a. 90%), primarily because of the designed conical intersection. A common characteristic among these dyes is their low quantum yield, as supported by photochemical measurements (Fig. 1C). A chloroform solution containing each PTD was prepared at a PTD molar ratio of 50:1. Following the thin-film method, chloroform solutions were cast and hydrated to prepare liposomes. After incorporation into the bilayers, the absorption peaks were observed at approximately 800 nm (Fig. 1D). Subsequently, solutions containing PTD-embedded liposomes were illuminated by an 808 nm laser (160 mW), resulting in a remarkable temperature increase in the bulk solution, except for the DDI dye (Fig. 1E). This might be attributed to the relatively low photostability and low incorporation of DDI compared with those in the others. No further size control was implemented, enabling straightforward observation of liposome behavior under an optical microscope (Fig. S2).

**Fig. 1.**
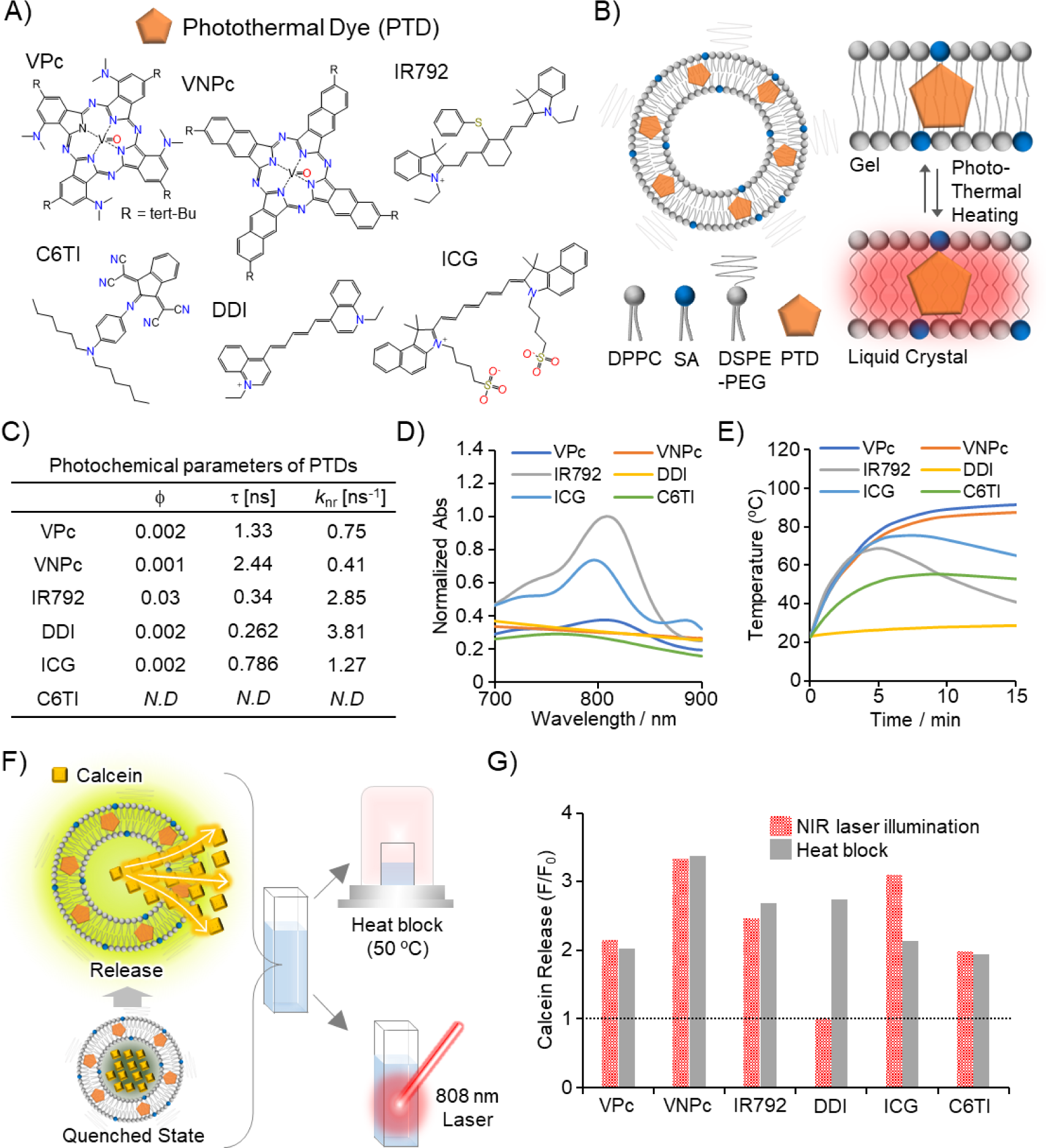
Screening of photothermal dyes (PTD) to develop near infrared (NIR)-modulated liposomes under bulk conditions. (**A**) Chemical structures of PTDs. (**B**) Schematic illustration of a PTD-embedded liposome for inducing phase transition upon NIR illumination. (**C**) Quantum yield (*φ*), fluorescence lifetime (*τ*), and rate constant of nonradiative relaxation (*k*_nr_) of each PTD. (**D**) Absorption spectra and (**E**) heat production ability of PTD-embedded liposomes induced by 808 nm laser illumination (160 mW). (**F**,**G**) Analysis of the heat-induced leakage of calcein from PTD-embedded liposomes. Each PTD-embedded liposome suspension was heated on a heat block at 50 °C for 20 min or illuminated by an 808 nm laser (160 mW) for 5 min. The changes in the normalized fluorescence intensity of calcein (F/F_0_) were evaluated in both heating methods (G). The dotted line represents a control line without heating treatments (F/F_0_ = 1 indicates no calcein leakage).

To investigate phase transition behavior, we prepared PTD-embedded liposomes encapsulating a high concentration of calcein as a hydrophilic payload (50 mM) (27). Specifically, when a phase transition occurs and calcein leaks, the fluorescence-quenched state of calcein is resolved, resulting in enhanced fluorescence. This enhanced fluorescence served as an indicator of bilayer loosening. To validate the evaluation system, we studied the release of calcein from liposomes by heating them on a heat block (50 °C, 20 min) (Fig. 1F). In all cases, fluorescence enhancement was observed after heating, suggesting that PTD did not interfere with the inherent phase transition properties of the bilayers (Fig. 1G). Instead of the heat block, when an 808 nm laser illuminated the liposome solution, the temperature was elevated above *T*_c_, and calcein release was also observed for all PTDs except DDI, which was consistent with the results shown in Fig. 1E. These findings indicate that a phase transition is induced when the medium temperature surpasses the threshold. Despite the differences in photothermal properties and incorporation efficiency among the PTDs, few considerable variations in calcein release were detected under bulk conditions, at least in this context.

### Microscopic observation of PTD-embedded liposomes

We further investigated the microscopic-scale behavior of the phase transition triggered by NIR laser illumination. Similar to the evaluation conducted under bulk conditions, we assessed the occurrence of phase transition by monitoring the fluorescence enhancement resulting from calcein leakage. A solution of PTD-embedded liposomes encapsulating calcein was cast onto a glass-bottomed dish for observation at the micron scale. To enable real-time visualization of calcein leakage during photothermal heating with NIR laser illumination, we employed a confocal microscopic system equipped with an 808 nm laser (Fig. 2A) (28). All microscopic experiments in the present study were conducted at 30 °C. Consequently, when the VPc-embedded liposomes (hereafter VPc-Lipo) were stimulated by an 808nm laser (10 s, 950 μW) during microscopic observation, the fluorescence intensity of calcein increased (Fig. 2B, upper panel). Once NIR illumination ceased, the intensity returned to basal levels, indicating that bilayer loosening was not irreversible and the leakage event was controllable. Similarly, we tested other PTD liposomes but rarely observed significant fluorescence changes upon NIR illumination [Fig. 2B (lower panel), Fig. 2C, D]. Notably, in contrast to the results obtained under bulk conditions, only VPc-Lipo exhibited calcein leakage, suggesting the induction of an effective phase transition in the bilayer at the microscopic scale (Fig. 2 B–D, Fig. S3).

**Fig. 2.**
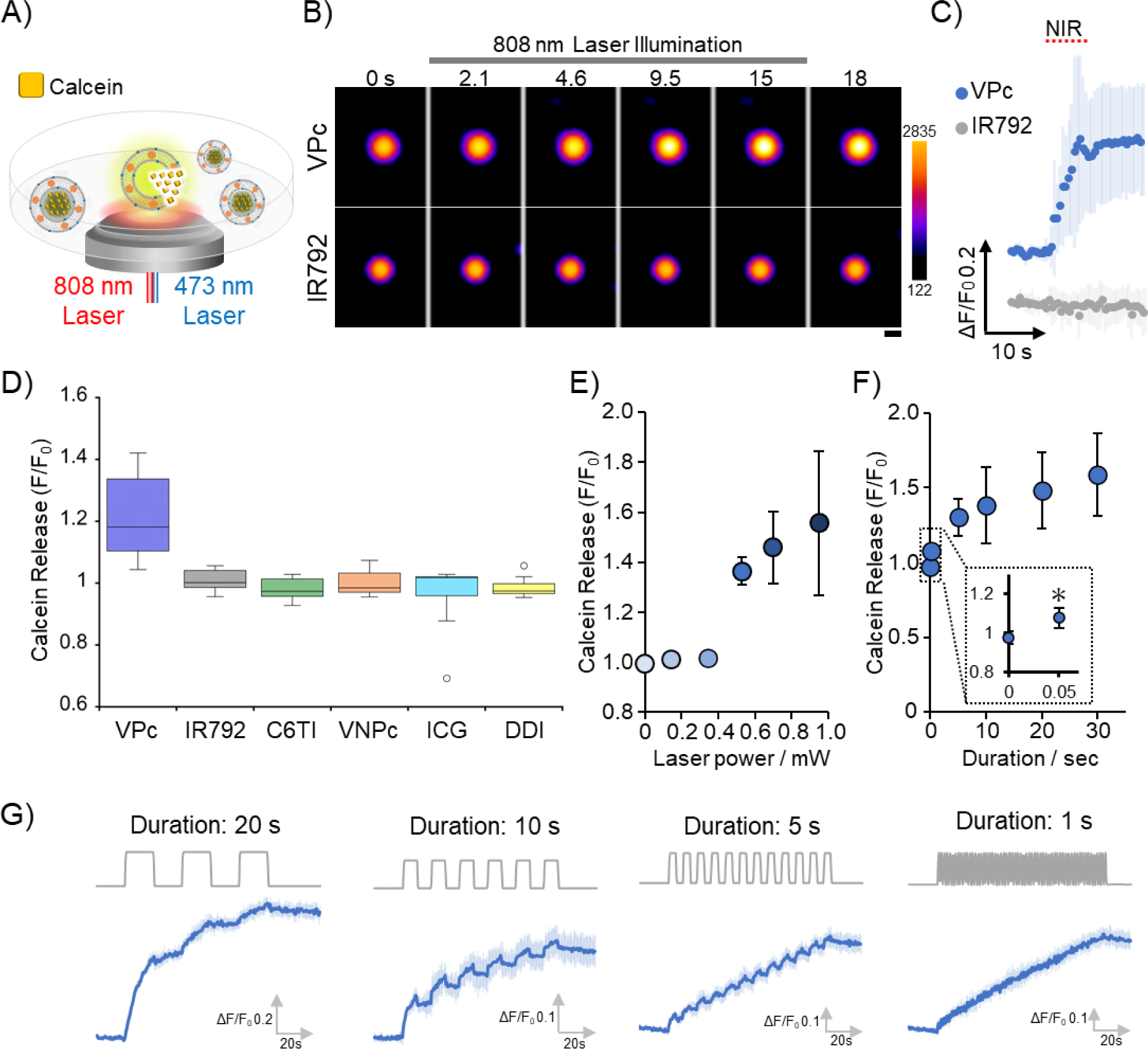
Calcein release from photothermal dye (PTD)-embedded liposomes at the microscopic level. (**A**) Microscopic setup to examine calcein leakage upon 808 nm laser illumination. Fluorescence changes were observed using a 473 nm laser. (**B**) Time course montage of calcein leakage from VPc- or IR792 embedded liposomes (upper panel: VPc, lower panel: IR792). Scale bar: 2 μm. (**C**) The mean of calcein release (F/F_0_) ± standard deviation (SD) (*n* = 8–10) was plotted in the time course. The red dashed line represents the timing of 808 nm laser illumination. (**D**) Evaluation of calcein release from PTD-embedded liposomes following 808 nm laser illumination (10 s, 950 μW). The line in the boxes shows the median (*n* = 10). (**E**) Calcein release from VPc-embedded liposomes (VPc-Lipo) at different laser powers during 50 ms stimulation (mean ± SD, *n* = 5). (**F**) Calcein release from VPc-Lipo at different durations at a constant laser power of 950 μW (mean ± SD, *n* = 5). (**G**) Different temporal patterns of calcein release following 808 nm laser illumination (950 μW; 20, 10, 5, and 1 s). Each data point shows the mean ± SD (*n* = 5). VPc, phthalocyanine vanadium oxide complex

We further examined the properties of VPc-Lipo by varying laser power amplitude and duration. By varying laser power, we found that effective calcein leakage occurred above 530 μW (Fig. 2E). Next, we analyzed the leakage profile for varying time durations at a constant laser power of 530 μW. Owing to the microscopic setup, which allowed a maximum shutter speed of only 50 ms, the actual temporal resolution limit remains unclear. However, flash illumination (i.e., 50 ms) led to significant calcein leakage (Fig. 2F). In addition, NIR illumination was repeated for different durations (1, 5, 10, and 20 s) and resulted in a reversible response to illumination (Fig. 2G). Longer illumination times led to a rapid decrease in calcein within the liposomes through multiple leakage events, whereas shorter illumination periods resulted in a comparatively lower decline.

To gain further insights into the observed mechanism, we imaged membrane fluidity during photothermal heating using Flipper-TR as a fluorescence lifetime (FLIM)-based fluidity sensor (Fig. 3A) (32). The bilayer in VPc-Lipo was stained with Flipper-TR dye, and its fluorescence lifetimes were plotted against varying temperatures in the surrounding medium. As the change in fluidity gradually decreased with increasing temperature, the fluorescence lifetime decreased, reaching approximately 4.2 ns of Δτ_1_ at a threshold temperature corresponding to the phase transition from gel to liquid crystalline at approximately 41 °C (Fig. 3B). Similarly, membrane fluidity was analyzed upon illumination of VPc-Lipo with an 808 nm laser during microscopic observation. Notably, even when calcein leaked, the lifetime (τ_1_) remained 5.2 ns, which is not consistent with the value observed at the membrane when it reached 41 °C (Fig. 3C). The underlying mechanism is addressed in detail in the Discussion section. To further analyze membrane permeability during NIR illumination, we encapsulated fluorescein isothiocyanate (FITC)- conjugated dextran (FITC-Dex) of varying sizes (4, 40, and 500 kDa) within the liposomes. Relatively large FITC-Dex molecules, such as those of 4 and 40 kDa, leaked out, whereas 500-kDa FITC-Dex was retained inside the liposome (Fig. 3D). These results, showing that Dex molecules with relatively large molecular weight leaked, indicate that phase transition is associated with bilayer structure cracking and increased membrane fluidity (33).

**Fig. 3.**
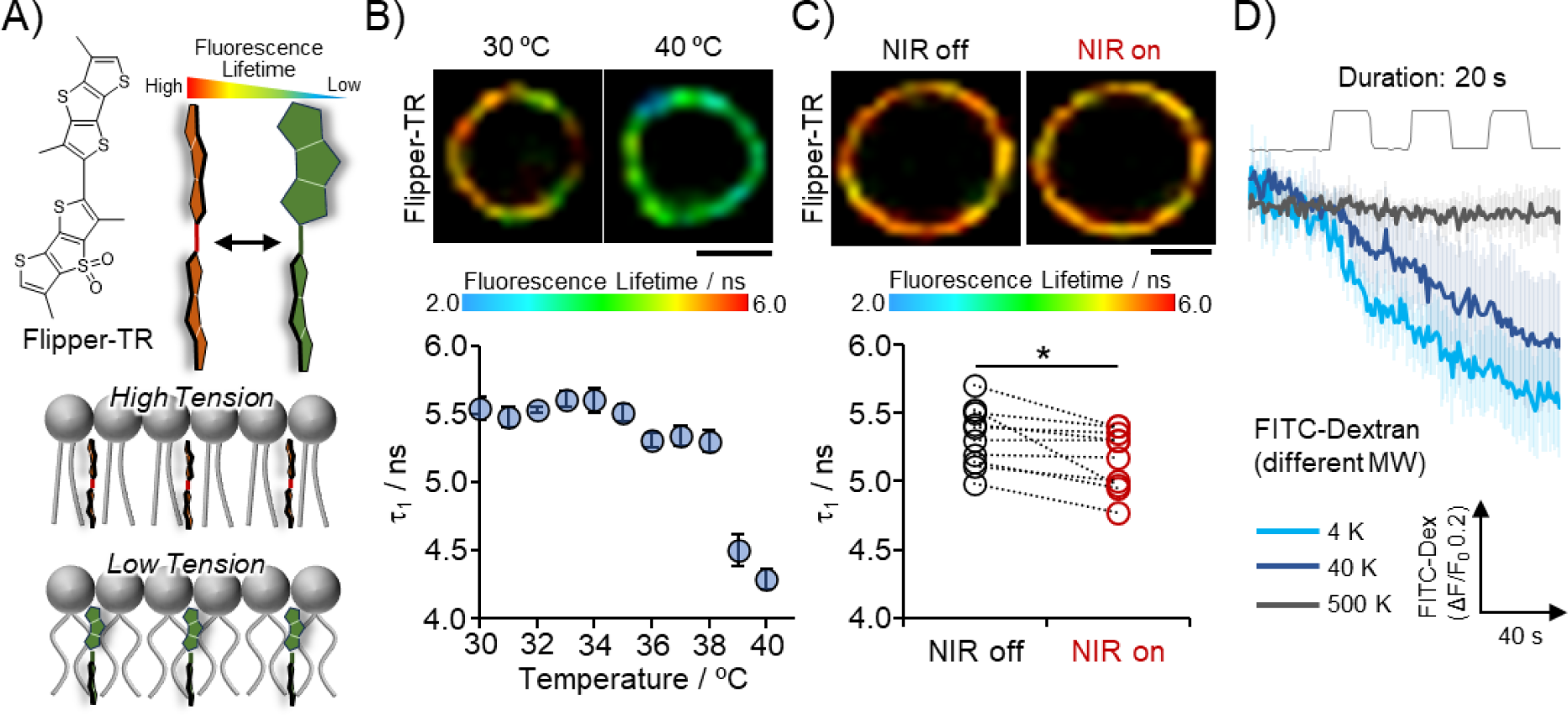
Minimal alteration of membrane fluidity in VPc-embedded liposomes following illumination with an 808 nm laser. (**A**) Schematic illustrations of Flipper-TR as a membrane tension FLIM sensor. (**B**) Calibration curve of Flipper-TR against temperature. The fluorescence lifetime (*τ_1_*) of Flipper-TR, which was located at the bilayer in VPc-Lipo, was plotted against different temperatures (*n* = 11). Scale bar: 3 μm. (**C**) Visualization of membrane tension upon 808 nm laser illumination (950 μW) (NIR off: 5.3 ± 0.21 ns, NIR on: 5.2 ± 0.22 ns, *n* = 10). Paired *t*-test, \**p* < 0.05. Scale bar: 5 μm. (**D**) Bilayer permeability following 808 nm laser illumination (950 μW). The leakage of encapsulated FITC-labeled dextran of different molecular weights (4k, 40k, and 500k) was analyzed (each data point shows mean ± SD, *n* = 4–5). VPc, phthalocyanine vanadium oxide complex; FLIM, fluorescence lifetime imaging microscopy.

### Evaluation of temperature increment in proximity to lipid bilayers

We sought to examine whether a PTD could effectively induce local heating within the bilayers, i.e., “thermal confinement” in the membranes of VPc-Lipo. Since heat dissipation and transfer are nonequilibrium and short-lived processes, capturing these events using conventional optical microscopes is challenging. However, if the temporal resolution is compromised, fluorescence thermometry, which is capable of sensing temperature changes through detectable fluorescence signals, such as fluorescence intensity and lifetime, is practical for approximating the temperature increment as an equilibrium state. Recently, we established a method that enables microscopic-scale thermometry with a temperature-sensitive dye using fluorescence lifetime imaging (FLIM) microscopy (34, 35). Regarding accuracy, FLIM-based analysis is superior to intensity-based analysis. In the current study, tetramethylrhodamine isocyanate (TRITC), a typical temperature-sensitive dye, was used for FLIM thermometry close to the bilayer. As direct thermometry inside the bilayers is challenging under highly fluidic conditions, we positioned the temperature sensor close to the bilayer rather than embedding it directly into the membranes (36). To achieve this, TRITC was placed a few nanometers away from the membrane surface, where TRITC-labeled avidin was anchored to the bilayers through a PEG linker with biotin at the terminus (Fig. 4A). Importantly, although the size of the liposomes ranged from 50 nm to a few micrometers, unilamellarity was observed in most cases (Fig. S2). Cryo-TEM images clearly indicated that avidin was located near the membrane (Fig. 4A). Before the experiments, the fluorescence lifetime of TRITC was plotted against temperature to generate a calibration curve (Fig. 4B). To validate this temperature-monitoring system further, we investigated the correlation between the detected temperature increase and calcein leakage. Magnetic particle agglomeration was conducted in a dish near the VPc-Lipo tethering TRITC. This setup enabled the creation of a localized heat source upon NIR illumination, subjecting the liposomes to external heating (Fig. 4C). By varying the amplitude of an 808 nm laser, calcein leakage was initiated above approximately 8 °C increment, as converted from the lifetime using the calibration curve (Fig. 4E, blue circle). Thermometry results in the cases where calcein leaked due to external heating indicated an average threshold temperature increment of approximately 10 ± 2.4 °C (Fig. 4F, Supporting Fig. S4). Previous studies on DPPC-based temperature-sensitive liposomes indicated that payload leakage was initiated at 37–38 °C and accelerated at 40 °C, consistent with our results(27). Similarly, we performed thermometry in close proximity to the bilayers when VPc-Lipo was directly exposed to NIR illumination, i.e., internal heating of the bilayer. At the lowest laser power of 530 μW, where calcein leakage was observed repeatedly as seen in Fig. 2E, the temperature increment was estimated to be 1.1 ± 1.4 °C from the calibration curve (Fig. S4, Fig. 4F). Moreover, as laser power increased to 950 μW, the fluorescence lifetime dropped, corresponding to a temperature change of 3.1 ± 2.5 °C (Fig. S4, Fig. 4F). Notably, when membrane loosening occurred during both external and direct local heating, distinct temperature gradients were formed on the lipid bilayers. In the case of local heating, the heat energy obtained from the photothermal effect seemed to be primarily directed toward altering bilayer dynamics. Consequently, the excess energy rarely dissipated, resulting in only a slight temperature increase a few nanometers from the surface.

**Fig. 4.**
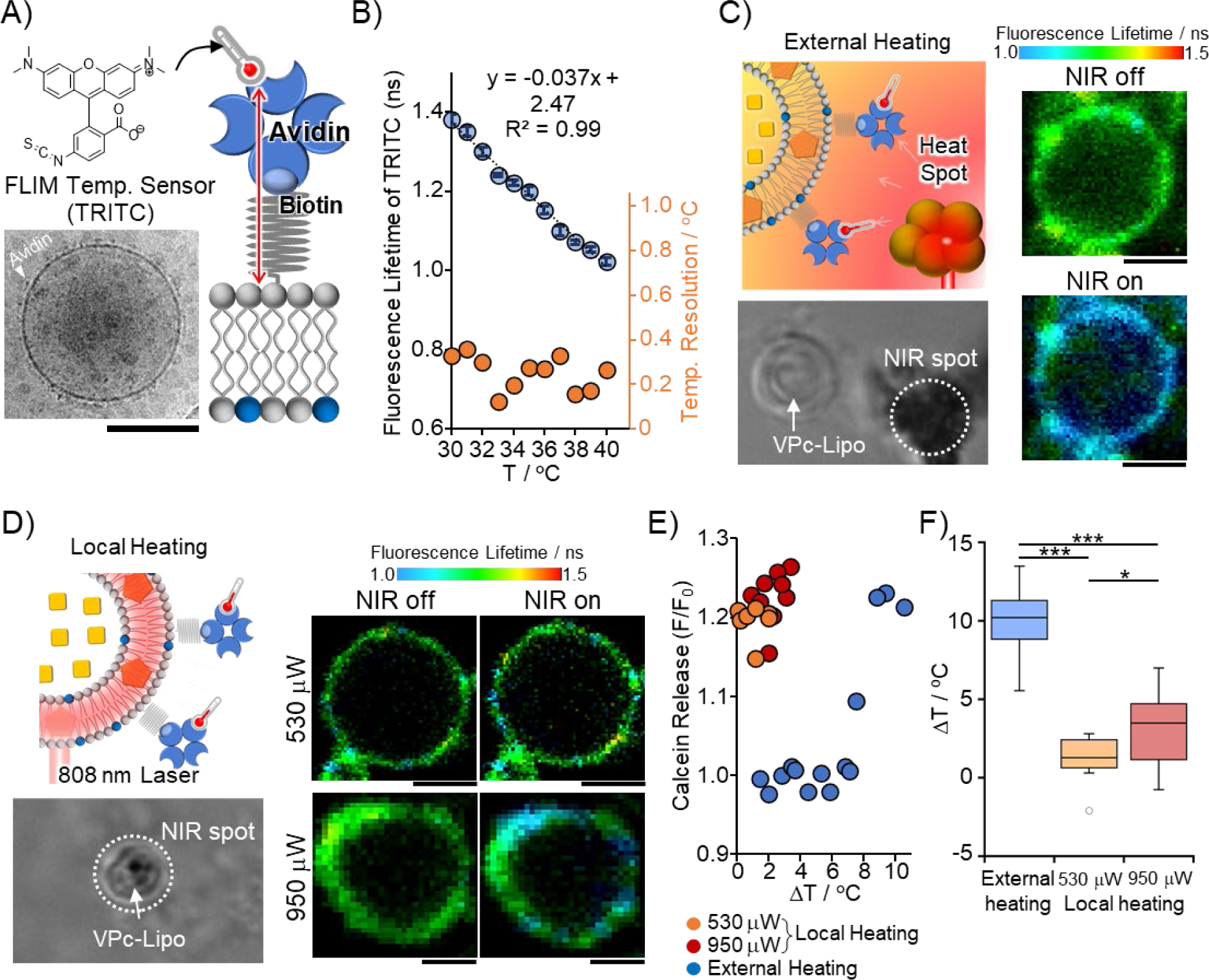
FLIM thermometry demonstrates efficient local heating near the lipid bilayers of VPc-Lipo. (**A**) Schematic illustration of FLIM thermometry near the VPc-Lipo bilayer using tetramethylrhodamine (TRITC) as a FLIM temperature sensor. Transmission electron microscopy of VPc-Lipo with avidin is shown. The white triangle indicates the representative avidin structure. Scale bar: 100 nm. (**B**) Calibration curve of TRITC against various temperatures (mean ± SD, *n* = 11). The secondary axis denotes the accuracy of thermometry derived from the SD value at each datapoint (*y* = -0.037*x* + 1.37, *R*^2^ = 0.99). (**C**) FLIM thermometry at the bilayer surface via external heating. Magnetic particles (Sicaster-M-CT) act as photothermal material in the glass-bottomed dish, generating external heat. FLIM images with and without 808 nm laser illumination (950 μW). Scale bar: 2 μm. (**D**) FLIM thermometry at the bilayer surface following local heating (808 nm laser: 530 and 950 μW). Scale bar: 2 μm. (**E**) Fluorescence change (F/F_0_) of calcein release against temperature increments (ΔT) near the bilayer. ΔT was estimated from the calibration curve (**B**) for external and local heating (530 and 950 μW, respectively). (**F**) Assessment of temperature increments near the bilayer under successful calcein release conditions. The line within the box represents the median. Statistical analysis was conducted using the Holm method (*n* = 10). Significance denoted as \*\*\**p*< 0.0001, \**p*< 0.05. Mean ± SD values indicate 10 ± 2.4 °C (external heating), 1.1 ± 1.4 °C (530 μW local heating), and 3.1 ± 2.5 °C (950 μW local heating). VPc, phthalocyanine vanadium oxide complex; FLIM, fluorescence lifetime imaging microscopy.

### Molecular dynamics simulation

As a reference, differential scanning calorimetry was used to analyze the extrapolated onset temperatures and full width at half maximum of the phase transition in PTD-embedded liposomes (Fig. 5A, Fig. S5). Although PTDs did not drastically alter phase transition, significant differences compared with liposomes without dyes were observed in the particular cases of VPc and IR792. Using molecular dynamics (MD) simulations, we further investigated the mechanism by which VPc allows for effective local heating due to thermal confinement in lipid membranes. Three selected dyes were placed in water, at the water-lipid membrane interface, and inside the lipid membrane, and relaxed by equilibrium MD at 1 atm and 300 K in an NPT ensemble for 5 ns, followed by estimation of the stabilization energy of each dye (Fig. 5B). After the energy of the dyes in water was standardized, the stabilization energies were evaluated for each environment. While IR792 and C6TI at the water-lipid membrane surface were more stable than those inside the lipid membrane, VPc favored the internal space of the membrane, indicating a high affinity for lipid bilayers (Fig. 5C). The generated heat was likely effectively transferred to the membrane when the dye was incorporated into the lipid interior. Furthermore, to investigate heat propagation through lipid membranes in the simulations, VPc and IR792 were placed at favorable positions, i.e., inside the lipid membranes and on the surface of the membrane, respectively. Subsequently, the thermal relaxation structure of dye-lipid complexes was constructed using equilibrium MD (Fig. 5D). Assuming that the dye underwent photothermal conversion, resulting in a temperature rise and stabilization at 350 K in nonequilibrium MD, we evaluated heat propagation by estimating the time constant for the lipid membrane to reach 350 K. As a result, the heat propagation time constant, defined as τ, for VPc and IR792 was estimated to be 3.83 ns in the membrane and 4.64 ns at the interface, respectively (Fig. 5E,F). It can be reasoned that membrane motion could directly consume the heat produced by VPc. In contrast, as IR792 is located at the interface of the membrane, heat might be partially dissipated to the water, resulting in a less effective perturbation of the membrane. As another possible mechanism, given that IR792 is also partially located inside the lipid membrane, the dye strongly interacted with the hydrophobic part of the lipid, resulting in a boundary in the ordered layer with an aligned lipid orientation and a complex, layered structure different from that of a normal lipid bilayer (Fig. S6). This boundary impedes the heat propagation path by atomic vibration, resulting in a low heat propagation time constant (τ = 5.42 ns) (Fig. 5F).

**Fig. 5.**
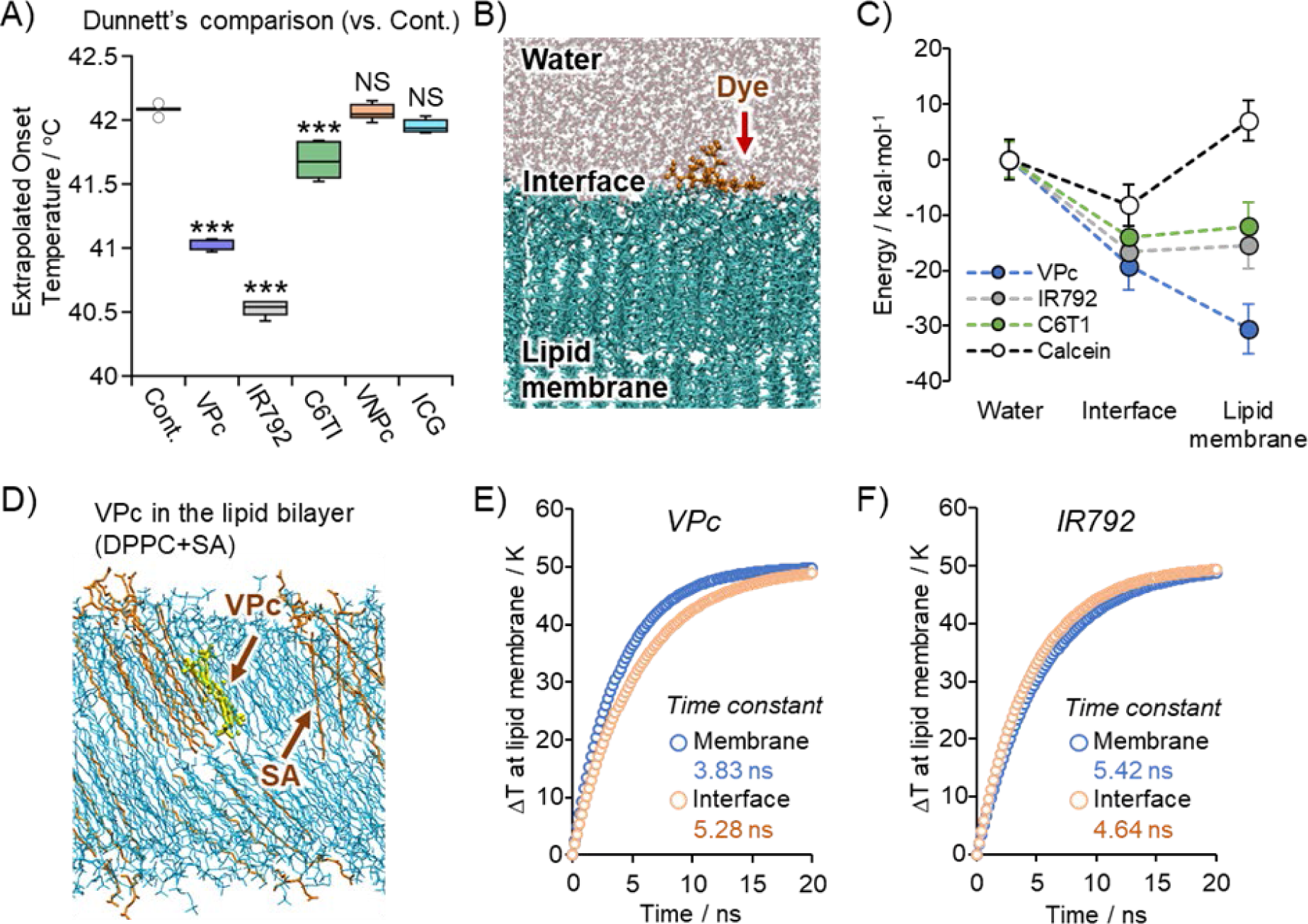
Stabilized energy of photothermal dye (PTD) within the lipid bilayers and heat propagation through molecular dynamics simulation. (**A**) Extrapolated onset temperatures were measured by differential scanning calorimetry (*n* = 6) using the same lipid formulation as that in Fig. 1. Each data point represents the mean ± SD. Dunnett’s multiple comparison test indicated significant differences (\*\*\**p* < 0.0001). (**B**) Lipids (DPPC and SA) and water molecules are depicted in green and gray, respectively. (**C**) The potential energy of each PTD and calcein was evaluated relative to their energy in water, with data variation derived from each time step (0.5 ps) over 1 ns. (**D**) Stability analysis of VPc in a lipid membrane composed of DPPC and SA. (**E**,**F**) Heat propagation in lipid bilayers via local heating with VPc (**E**) or IR792 (**F**). Using nonequilibrium molecular dynamics, the time constant for the lipid membrane temperature to reach 350 K from a base temperature of 300 K was estimated, and ΔT at the lipid bilayers was plotted on the *y*-axis. DPPC, 1,2-dipalmitoyl-sn-glycero-3-phosphocholine; SA, L-glutamic acid, *N*-(3-carboxy-1-oxopropyl)-,1,5-dihexadecyl ester; VPc, phthalocyanine vanadium oxide complex.

### Generation of the acetylcholine concentration jump using NIR-modulated liposomes

We further demonstrated the considerable potential of distinct local heating with thermal confinement as a platform for generating microscopic tools. Our proposed concept focused on encapsulating a bioactive compound as payload inside VPc-Lipo instead of a hydrophilic calcein dye. We anticipated that this liposome would allow on-demand release of the encapsulated payload at specific target locations upon NIR illumination. This concept is distinguished by the effective confinement of heat generated during NIR illumination within the lipid membrane, ensuring minimal impact on target biospecimens, such as tissues and cells. To this end, as shown in Fig. 6A, we prepared VPc-Lipo, in which avidin was immobilized on the surface through a PEG linker. Additionally, biotinylated iRGD peptides to avidin facilitated the anchoring of liposomes to live cell membranes (Fig. 6A). To assess the extent to which live cells received heat delivered from VPc-Lipo, we employed FLIM-based fluorescent thermometers targeting the plasma membrane (PTG) and endoplasmic reticulum (ER) (ETG) in HeLa cells (Fig. 6B, D) (34). As a result, FLIM thermometry using PTG and ETG estimated temperature increments of 0.39 ± 1.6 °C and 0.88 ± 0.43 °C, respectively, at 950 μW of the 808 nm laser (Fig. 6C,E,F). This liposomal system delivered the encapsulated payload with negligible thermal damage to the biospecimens, demonstrating the potential of NIR-modulated liposomes.

**Fig. 6.**
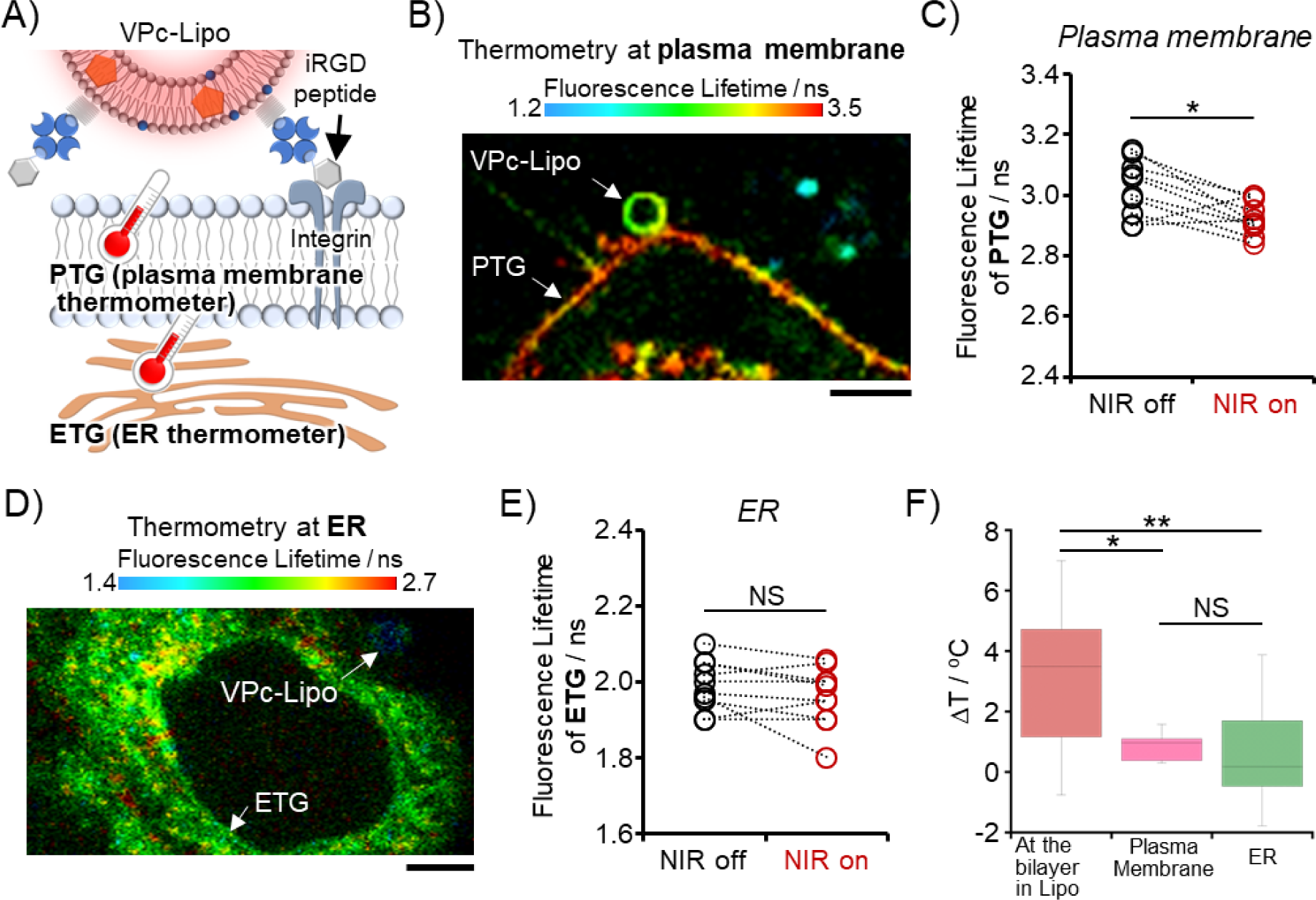
Minimal thermal damage following local heating as revealed by FLIM cellular thermometry. (**A**) Schematic illustration of VPc-Lipo anchored to the cellular membrane via interaction with the iRGD peptide and integrin. Temperature increments during the local heating of VPc-Lipo were analyzed using FLIM-based organelle thermometers [plasma membrane: PTG and endoplasmic reticulum (ER): ETG]. The lipid of VPc-Lipo was stained with DiI dye. (**B**) FLIM image of PTG in HeLa cells with VPc-Lipo. Scale bar: 20 μm. (**C**) Temperature increments at the plasma membrane (PTG) with NIR off/on (808 nm laser, 950 μW) (*n* = 10). Paired *t*-test, \**p* < 0.05. (**D**) FLIM image of ETG in HeLa cells with VPc-Lipo. Scale bar: 10 μm. (**E**) Temperature increment at ER (ETG) with NIR off/on (808 nm laser, 950 μW) (*n* = 10). Paired *t*-test, not significance (NS). (**F**) Temperature increments estimated from the calibration curves. Mean ± SD represents 3.2 ± 2.4 °C at the lipid bilayer (950 μW), 0.39 ± 1.6 °C at the plasma membrane, and 0.88 ± 0.43 °C at ER. Tukey’s multiple comparison indicates significance (*n* = 10). \**p* < 0.05, \*\**p* < 0.01. VPc, phthalocyanine vanadium oxide complex; FLIM, fluorescence lifetime imaging microscopy.

Subsequently, we encapsulated acetylcholine (ACh) within these liposomes and tethered the iRGD peptide (Fig. 7A). ACh was maintained at concentrations in the order of hundreds of millimolars with reference to natural synaptic vesicles (37). Neuronal and muscular systems, where neurotransmitters play pivotal roles, are highly susceptible to thermal stress and the subsequent induction of Ca^2+^ flux (38, 39). Thus, our liposome system, characterized by thermal confinement, fulfills the requirements for modulating the spatiotemporal dynamics of neurotransmitters in these delicate biospecimens. To visualize the changes in ACh concentration, GACh2.0, a fluorescent sensor for detecting ACh, was expressed on the surface of HEK293T cells (Fig. 7B) (40). The normalized intensities of GACh2.0 were plotted at different concentrations to generate a calibration curve (Fig. 7C). When liposomes were placed on the cellular membrane and subjected to NIR illumination, we observed a rapid increase in ACh levels at the moment of illumination, resulting in the generation of a concentration gradient across a few cells (Fig. 7D, E). The concentration of ACh, estimated from the calibration curve, could reach several hundred micromolars at the closest region of interest (ROI) (Fig. 7C–F). Additionally, we confirmed that a brief 50 ms stimulation was sufficient to release ACh a few times sequentially (Fig. 7G–H). Notably, liposomes of different sizes produced various spatial ACh concentration patterns. The prepared suspension contained a small-sized liposome of approximately 100 nm (Fig. S2). When a small-sized liposome was on the cellular membrane, the spatial distribution of ACh generated using NIR illumination was confined to a single cell (Fig. 7J–K). Although it has not been systematically investigated, size control of liposomes could potentially offer a more precise means for modulating spatial distribution in future applications.

**Fig. 7.**
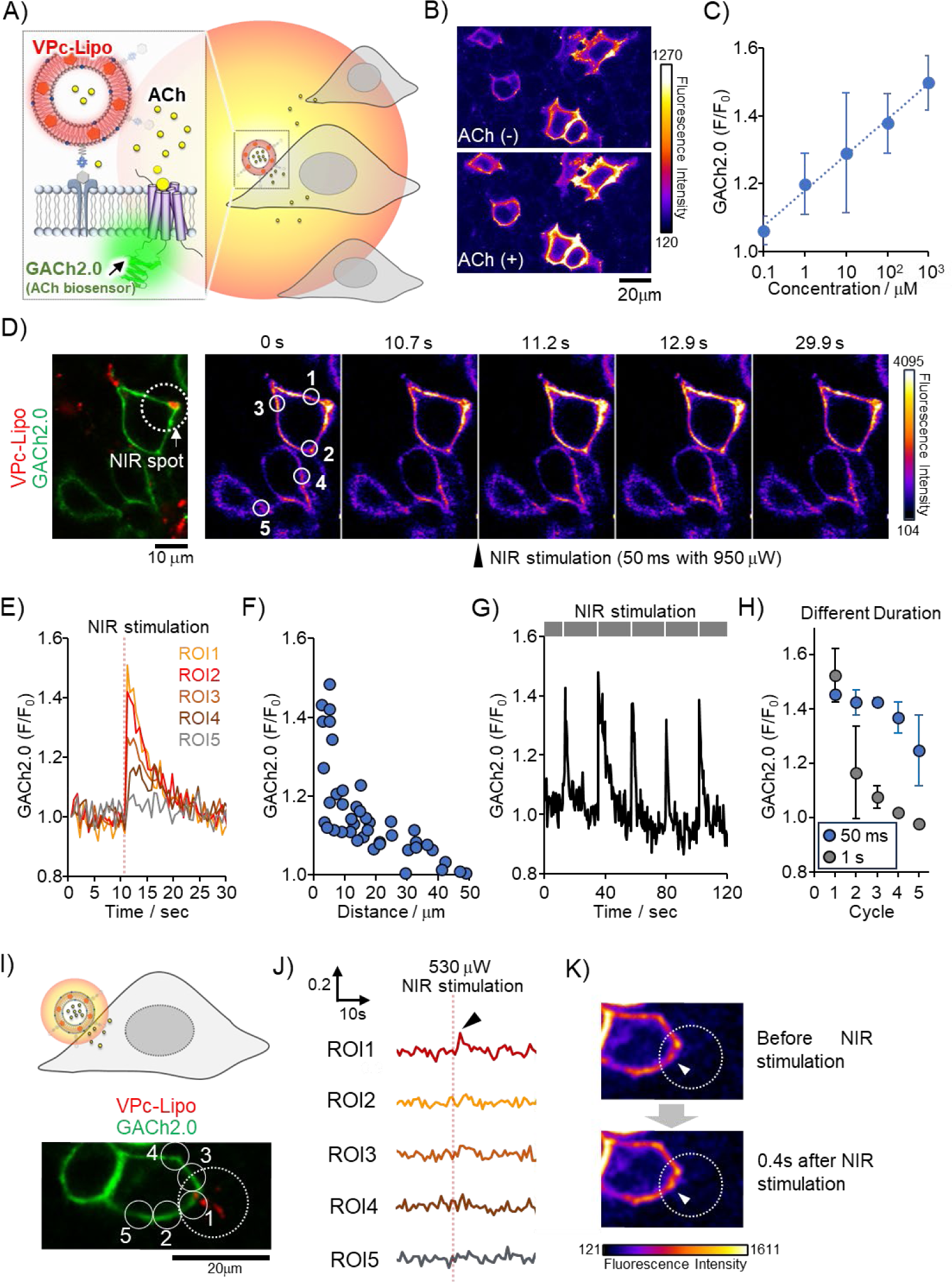
Validation of the release profile of acetylcholine using near infrared (NIR)-modulated VPc-Lipo. (**A**) Schematic illustration of VPc-Lipo encapsulating acetylcholine (ACh) anchored to the cellular surface, followed by 808 nm laser illumination. Spatiotemporal dynamics of ACh visualized by GACh2.0 as an ACh biosensor. (**B**) Fluorescence images of GACh2.0 in HEK293T cells in the absence and presence of 100 mM ACh. Scale bar: 20 μm. (D) Calibration curve correlating the normalized fluorescence intensity (F/F_0_) of GACh2.0 with ACh concentration (μM) (*y* = 0.0458ln(*x*) + 1.18, *R*^2^ = 0.993). Each plot indicates mean ± SD. (E) Fluorescence images of VPc-Lipo (red) in HEK293T cells expressing GACh2.0 biosensor (green). Montage of ACh release from VPc-Lipo upon 808 nm laser illumination (50 ms, 950 μW). Scale bar: 10 μm. (**E**) Analysis of the normalized F/F_0_ of GACh2.0 at different regions of interest (ROI) over time [relevant to (**D**)]. (**F**) Gradient of ACh concentration in the observation area under the microscope. Normalized F/F_0_ plotted against distances from the NIR spot. Distance on the *x*-axis is defined as the distance from VPc-Lipo to ROI (*n* = 45). (**G**) Fluorescence responses of GACh2.0 upon sequential illumination using an 808 nm laser (five times, 50 ms). (**H**) Gradual decrease of normalized F/F_0_ of GACh2.0 following five times illumination (mean ± SD, *n* = 3). (**I**) Schematic illustration and fluorescence image of small-sized VPc-Lipo (red) in HEK293T cells expressing GACh2.0 biosensor (green). (**J**) Time course analysis of the normalized F/F_0_ of GACh2.0 at different ROIs within the same cell (HEK293T). Stimulation was conducted at 530 μW for 50 ms. (**K**) Visualization of local concentration jump of ACh [same dataset as (**I**–**J**)]. VPc, phthalocyanine vanadium oxide complex.

### Modulation of neuronal functions in skeletal muscle and the *Drosophila* brain

Finally, we explored the applications of the liposomal system in skeletal muscles and the brain of *Drosophila*, both of which express ACh receptors on their cell membranes. As a model for skeletal muscles, we stained C2C12 myotubes with a fluorescent Ca^2+^ indicator (Fluo-4 AM) (Fig. 8A). Upon NIR illumination of the liposomes adhering to the myotubes, we observed an elevation in cytoplasmic Ca^2+^ levels in response to the release of ACh from the liposomes (Fig. 8B). Moreover, sequential NIR illumination led to the release of ACh, which subsequently induced Ca^2+^ elevation (Fig. 8B,C). The Ca^2+^ response in the 5^th^ illumination cycle was smaller than that in the previous cycles, suggesting that ACh release eventually declined, consistent with the results shown in Fig. 7H. This elevation was also attributed to the stimulation of ACh receptors, as evidenced by the limited cytoplasmic Ca^2+^ alteration in the presence of α-bungarotoxin, an Ach receptor inhibitor, during NIR illumination (Fig. 8D).

**Fig. 8.**
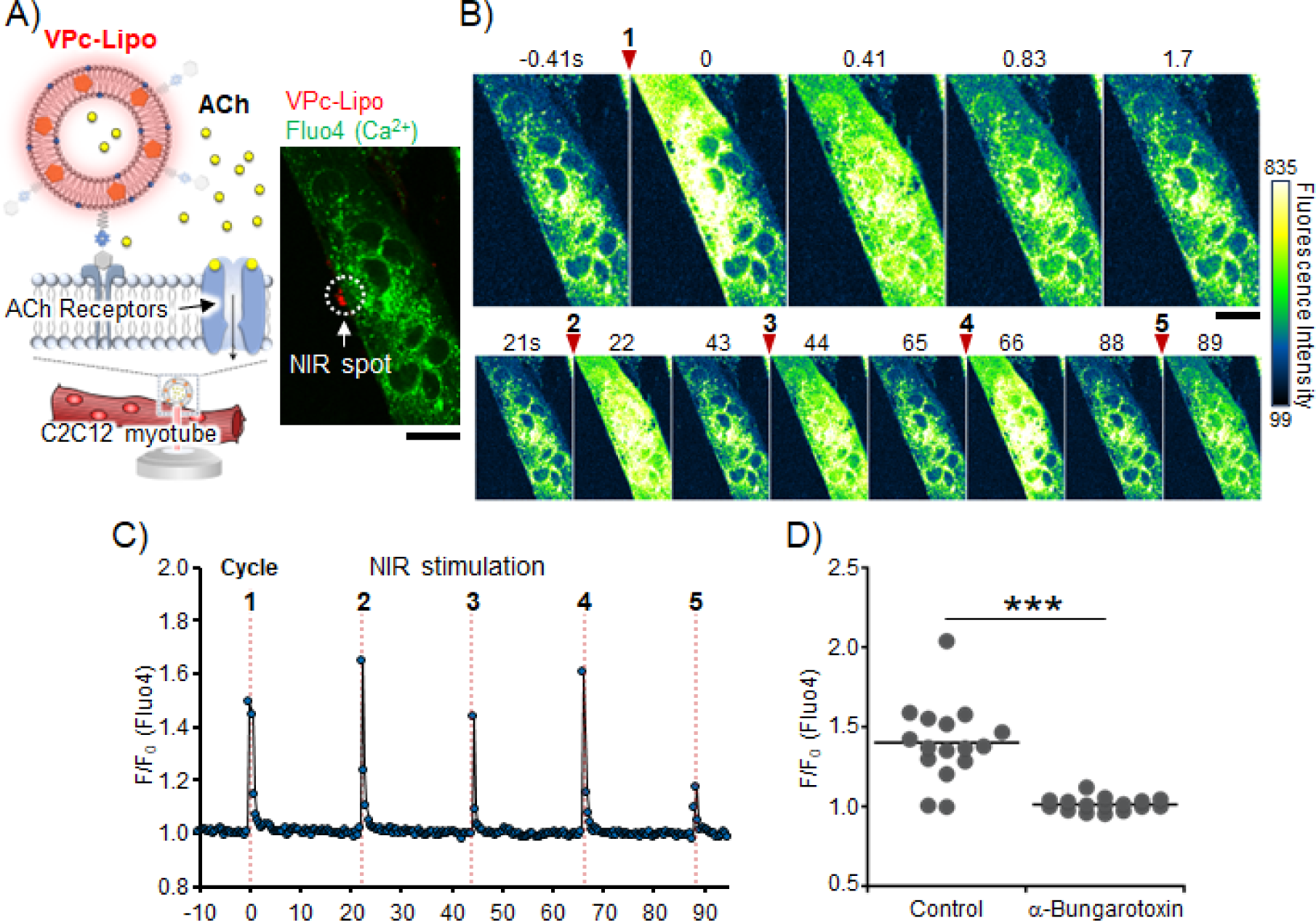
Ca^2+^ imaging in C2C12 myotubes induced by near infrared-modulated ACh release from VPc-Lipo. (**A**) Schematic illustration of VPc-Lipo encapsulating acetylcholine (ACh) anchored to the surface of skeletal muscle (C2C12 myotubes), followed by 808 nm laser illumination. Fluorescence images of Ca^2+^ sensor (Fluo-4) and VPc-Lipo (red) in C2C12 myotubes. Scale bar: 20 μm. (**B**) Montage of Ca^2+^ elevation induced by ACh release from VPc-Lipo upon 808 nm laser illumination in myotubes (five times, 50 ms, 950 μW). Scale bar: 20 μm. (**C**) Time course analysis of F/F_0_ of Fluo-4 in a single cell. (**D**) Inhibition experiments with α-bungarotoxin, an ACh receptor inhibitor (5 μM). The peak of normalized F/F_0_ upon 808 nm laser illumination [in the first cycle as shown in (**C**)] was plotted (region of interest number = 16, cell number = 8, independent 4–6 dishes). Student’s *t*-test indicated significant differences (\*\*\**p* < 0.0001). VPc, phthalocyanine vanadium oxide complex.

We extended this system to *Drosophila* brains, which express jGCaMP7c as a Ca^2+^ biosensor (41). Similarly, we introduced liposomes to the brain samples (Fig. 9A), followed by the illumination of VPc-Lipo with an 808 nm laser (50 ms, 950 μW) targeting the liposomes. The fluorescence intensity of jGCaMP7c in a single cell close to the NIR spot increased in response to the release of ACh from liposomes and returned to basal levels over several seconds (Fig. 9B). When relatively large liposomes were illuminated, the concentration gradient of ACh likely reached several cells surrounding the NIR spot (Fig. 9C). Notably, individual cells behaved differently in response to repeated NIR illumination (Fig. 9D). This technique could be an excellent microscopic tool for identifying cell types or heterogeneity, even within the same cell type in neuronal cells. Inhibition experiments using α-bungarotoxin indicated that the changes in fluorescence were a result of ACh receptor stimulation (Fig. 9E). In addition, in the presence of donepezil, an ACh esterase inhibitor, the fluorescence intensity of jGCaMP7c increased continuously and did not return to basal levels (Fig. 9F). In the nervous system, esterase is known to quickly decompose ACh to maintain appropriate ACh levels. Thus, esterase inhibition caused a continuous increase in ACh.

**Fig. 9.**
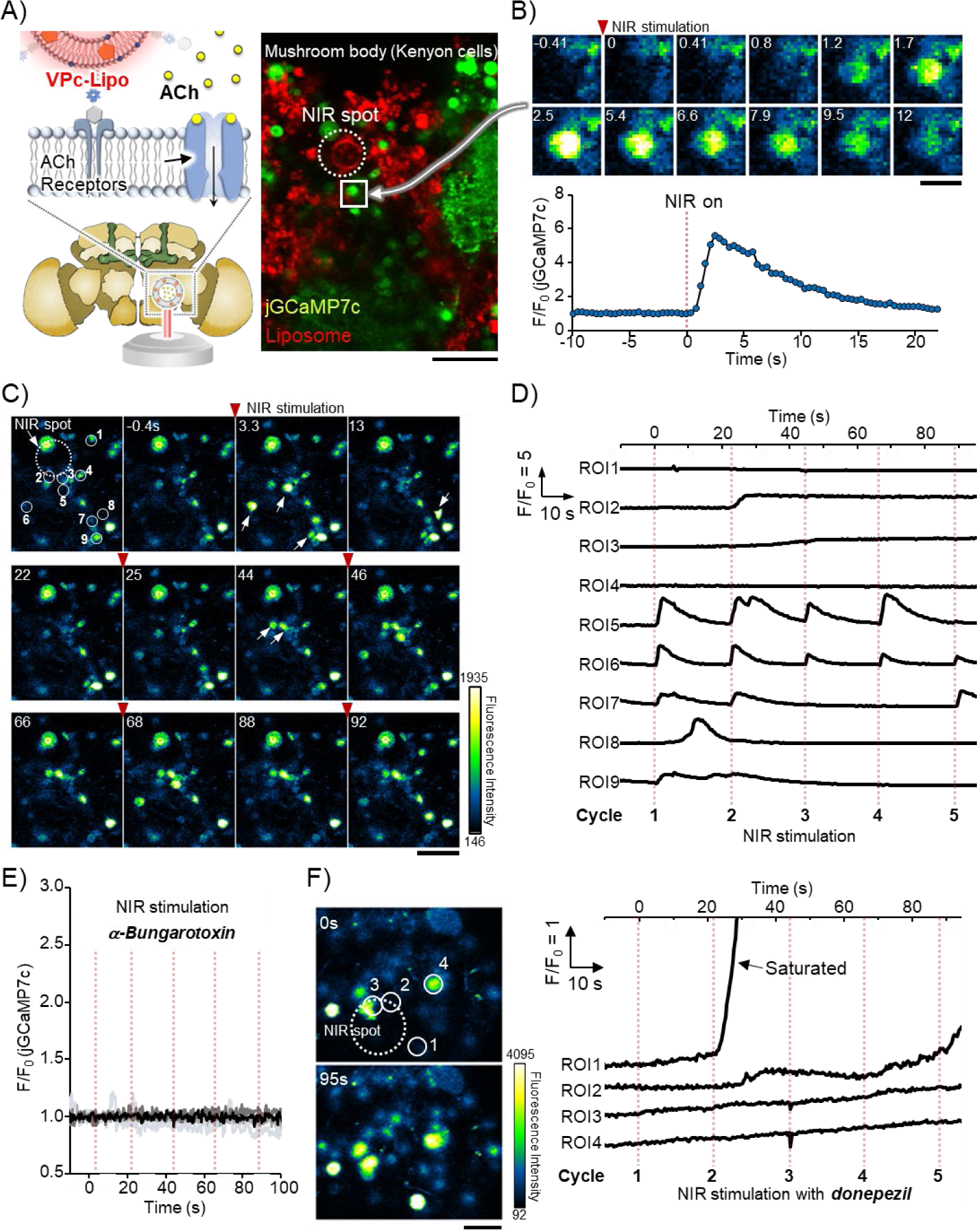
Imaging of acetylcholine (Ach)-evoked Ca^2+^ signals in *Drosophila* brain (*ex vivo*) using VPc-Lipo. (**A**) Schematic illustration of VPc-Lipo encapsulating ACh in the *Drosophila* brain (*ex vivo*), followed by 808 nm laser illumination. Fluorescence images of the brain expressing jGCaMP7c as a Ca^2+^ sensor (green) and VPc-Lipo (red). The white-lined square area indicates the target cell for imaging, as shown in (**B**). Scale bar: 20 μm. (**B**) Montage of Ca^2+^ dynamics in the cell (mushroom body) near VPc-Lipo. ACh was released from VPc-Lipo upon 808-nm laser illumination (50 ms, 950 μW). Scale bar: 5 μm. The normalized fluorescence intensity (F/F_0_) of jGCaMP7c was analyzed over time. (**C**, **D**) Montage of Ca^2+^ dynamics near the near infrared spot during 808 nm laser illumination (five cycles, 50 ms, 950 μW). Scale bar: 20 μm. Time courses were analyzed at different regions of interest (1–9). (**E**) Inhibition experiments with α-bungarotoxin, an ACh receptor inhibitor (5 μM). The thick line indicates the representative data (*n* = 8). (**F**) Inhibition experiments using donepezil as an ACh esterase inhibitor (100 nM). The experimental procedure was the same as that in (**C** and **D)**. Fluorescence images at the first illumination cycle (0 s) and 95 s (after the fifth illumination). Scale bar: 10 μm. VPc, phthalocyanine vanadium oxide complex.

## Discussion

In recent years, numerous studies have reported on photo-triggerable liposomes releasing their payloads through permeabilization of the lipid bilayers upon light illumination (25). For example, photo-crosslinking or light-induced oxidation of unsaturated lipids causes shrinkage or pore formation in the liposome, leading to payload release (26). In another pioneering study, light photoswitchable phosphatidylcholine with an azobenzene moiety perturbed lipid membranes mechanically during *cis*-*trans* isomerization, enabling the successful release of the drug (42). Although visible light illumination is unsuitable for biological applications, this issue can be circumvented using an upconversion particle that converts NIR light to visible light (43). Although these strategies appear robust, reversibility or the kinetic rate of release remains challenging. As mentioned earlier, local heating with thermal confinement in the lipid bilayers is one of the possibilities that allows for the fast release of the encapsulated payload while maintaining reversibility. In fact, 50 ms NIR illumination allowed ACh release from VPc-Lipo in the present study (Fig. 7G). However, this temporal resolution is constrained by the limited shutter speed in current confocal microscopy techniques. As the alteration of lipid membrane dynamics occurs at the microsecond scale, faster illumination should be feasible in the future. Similarly, the scanning rate in confocal microscopy settings limits the ability to capture the dynamics of ACh distribution. Although the generated steep ACh concentration gradient is evident, as shown in Fig. 7E,J, the actual local ACh concentration close to the lipid bilayers has not been properly evaluated owing to specific limitations.

Notably, in the context of local heating, a more recent study suggested that liposomes containing lipids tethering a porphyrin derivative as a photothermal agent could release calcein without a significant temperature rise in the medium (44). However, the associated heat dissipation and membrane fluidity alteration were not imaged. The present study demonstrates that local heating results in a negligible temperature increase near lipid bilayers. The change in membrane fluidity observed during NIR light illumination exhibited distinctive features compared with the fluidity change occurring uniformly across the lipid bilayers during external heating (Fig. 3C). Given the molar ratio of VPc to lipid (1:50), it is reasonable to infer that the phase transition events from a gel to the liquid crystalline state are likely confined to a very limited region, specifically the molecular environment surrounding the VPc dye at nanoscale levels. Consequently, this distinct photothermal heating created conditions under which the gel and liquid crystalline states coexisted within the bilayers. Previous studies suggested that such mixed-phase conditions result in grain boundary defects, which are highly effective in promoting the leakage of internal payloads (33). This hypothesis could also be supported by the fact that FITC-dextran of a few nanometers could leak out. The dynamics of membrane fluidity at the nanoscale level, which could not be captured in real-time owing to the spatial limitations of fluorescence microscopy, present an area for future research.

Our NIR-triggered liposomal system does not require lipid tethering of the dye, making it superior to previous studies. To investigate the importance of VPc, we synthesized VPc without the tert-butyl group, designated VPc-diMe, and examined calcein leakage capabilities and MD simulation (Fig. S7). We found that the VPc-diMe-embedded liposomes responded slightly to 808 nm laser illumination at a rate lower than that of VPc. MD simulation showed that VPc-diMe was less stable than VPc in lipid bilayers, whereas it was more stable than IR792 and C6TI (Fig. S8). This indicated that the tert-butyl substituent contributed to PTD dispersion in the hydrophobic region of the bilayers. Systematic studies of the structural relationships are required for a rational design that can work well in the future. As a microscopic tool for creating a concentration jump of bioactive compounds, the photocaging methodology has become a standardized technique. Once a photocleavable moiety initially masks the function of the compound, it can be recovered upon exposure to light. Despite being a robust technique, difficulties such as the limited photocaging compound availability, water solubility, and cleavage efficiency are frequently encountered. For instance, although ACh was discovered as the first neurotransmitter a century ago, photocaging has only been achieved recently due to its synthesis difficulty (45). While photocaging has been useful, solubility of photocaged ACh compound could reach approximately 50 μM, far from the physiological range of ACh in synaptic vesicles (45). In addition, uncaging efficiency is below 50% with byproduct formation. Therefore, alternative methods are in demand (46). Unlike photocaging methods, our liposomes mimic the concentration of ACh in actual synaptic vesicles and can release intact ACh without the photolysis of caged moieties. Another distinguishing feature is the use of NIR illumination, which is compatible with biospecimens, particularly for tissue and animal studies (Table S1) (47).

In conclusion, we successfully achieved thermal confinement in bilayers, as evidenced by FLIM thermometry and MD simulations. Our results will be useful for developing NIR-laser-operated biomaterials, where thermal damage is a critical concern. Furthermore, incorporating PTDs into liposomes is a valuable tool because of the simplicity of the lipid-dye mixture, which facilitates the construction of light-operated artificial synaptic vesicles. For instance, VPc-Lipo encapsulating ACh could find utility in unraveling diseases such as Alzheimer’s, where ACh esterase is highly expressed in the brains of patients (48). Additionally, the encapsulation of excitatory and inhibitory neurotransmitters such as glutamate and γ-aminobutyric acid can be extended to diverse applications in brain function modulation (49, 50). Moreover, encapsulating hydrophilic drugs, including mRNA and peptides, offers a pathway for more efficient drug delivery systems, particularly when activated using portable NIR laser devices (51).

## Materials and Methods

All chemicals were purchased from FUJIFILM Wako Pure Chemical Corporation (Osaka, Japan), Funakoshi (Tokyo, Japan), Merck (Darmstadt, Germany), TCI (Tokyo, Japan), Thermo Fisher Scientific (Waltham, MA, USA), Worthington Biochemical Corporation (Lakewood, NJ, USA), and YUKA SANGYO Co., Ltd. (Tokyo, Japan). Data were analyzed using Fiji (ImageJ; National Institutes of Health, Bethesda, MD, USA) and KaleidaGraph (Synergy Software, Reading, PA, USA).

### Preparation of PTD-embedded liposomes and iRGD-conjugated VPc-Lipo encapsulating ACh

A lipid mixture (5.0 mg) of 1,2-dipalmitoyl-sn-glycero-3-phosphocholine (DPPC), L-glutamic acid, *N*-(3-carboxy-1-oxopropyl)-,1,5-dihexadecyl ester (SA), and 1,2-distearoyl-sn-glycero-3-phosphoethanolamine-(polyethylene glycol) (DSPE-PEG) (9:1:0.06, molar ratio in chloroform, 250 μL) was prepared first. This was then mixed with each PTD stock solution (2 mM in chloroform, 62.5 μL) and DiD (2 mM in ethanol, 5 μL) in a glass vial. The molar ratio of DPPC to PTD was estimated to be 9:0.2. However, in the case of ICG, the dye ratio was changed to 9:0.02 because too much ICG drastically interfered with the phase-transition temperature. The solvent was evaporated to generate a thin film, which was then vacuumed to remove any residual solvent. The lipid film was hydrated in water (250 μL) containing 50 mM calcein, 200 mM sodium hydroxide, and 300 mM sodium gluconate at 60 °C for 10 min and vortexed immediately to form a suspension. After excess calcein was removed by size exclusion chromatography (PD10-column; Cytiva, Marlborough, MA, USA), PTD-embedded liposomes encapsulating calcein were obtained. To prepare tetramethylrhodamine (TRITC)-conjugated VPc-Lipo for FLIM thermometry, a liposome suspension was prepared according to the above-mentioned procedure using VPc dye and DSPE-PEG-biotin instead of DSPE-PEG. Subsequently, 10 μL of the liposome suspension and 2.5 μL of TRITC-labeled avidin stock solution (150 μM in water) were added to 150 μL phosphate-buffered saline (PBS) and incubated for 15 min. Excess TRITC-avidin that did not conjugate with the liposomes was removed by dialysis (Float-A-Lyzer®, MWCO: 300 kD; Funakoshi Co., Ltd., Tokyo, Japan) to obtain the liposome tethering TRITC-labeled avidin.

To prepare iRGD-conjugated VPc-Lipo encapsulating ACh, liposomes were prepared following the above-mentioned procedure using a lipid stock solution. DiD dye (2 mM in ethanol, 5 μL; PromoCell, Heidelberg, Germany) was added to the lipid solution to visualize the location of VPc-Lipo for microscopic studies. The thin lipid film was hydrated at 25 °C for 5 h in the presence of 500 mM ACh before incubating at 50 °C for 1 h and vortexing gently. Unencapsulated ACh and DiD dyes were removed using a PD-10 column with PBS as the eluent. Subsequently, the liposomes were first labeled with 40 µM avidin, followed by the removal of unconjugated avidin through dialysis using a Float-A-Lyzer® dialysis device. The avidin-labeled liposomes were conjugated with 0.08 µM iRGD-PEG-biotin. After removing unconjugated iRGD-PEG-biotin through dialysis using a Float-A-Lyzer® dialysis device, the finalized iRGD-conjugated VPc-Lipo encapsulating ACh was obtained. All liposomes were stored at 4 °C before experiments.

### Screening of PTD-embedded liposomes in bulk solution and under a microscope

Suspensions of the PTD-embedded liposomes were diluted with PBS (200×). The dispersion was heated using a heat block at 50 °C for 20 min or illuminated using an 808 nm NIR laser for 5 min. Calcein release was examined at 530 nm (excitation at 480 nm) using a Spectra Max Gemini XPS plate reader (Molecular Devices, San Jose, CA, USA). For microscopic evaluation, a liposome suspension encapsulating calcein (5 μL) was diluted with PBS (195 μL). The dispersion was then dropped onto a glass-bottomed dish and incubated overnight. Fluorescence images of the leakage of calcein from liposomes were captured at 256 × 256 pixels every 0.56 s using a confocal microscope (FV1200; Olympus Corporation, Tokyo, Japan) equipped with a 60× oil immersion objective lens (NA = 1.42, PLAPON; Olympus Corporation). The emission was accumulated in the FITC channel (emission filter set: 490–540 nm; laser: 473 nm). During observation, liposomes were illuminated with an 808 nm laser (530–950 μW, 0.05–10 s, laser core size: 50 μm; FiberLabs Inc., Saitama, Japan).

### FLIM analysis of temperature increment on the surface of liposomes and organelles

FLIM images were captured at 512 × 512 pixels every 1.66 s using a confocal microscope equipped with a 560 nm pulsed laser and rapid-FLIMHiRes with a MltiHarp 150 Time-Correlated Single Photon Counting unit (PicoQuant, Berlin, Germany). The TRITC emissions were accumulated using a filter (600/50 Bright Line HC; Semrock, Rochester, NY, USA). All fluorescence lifetime data were analyzed using SymPhoTime 64 (PicoQuant). The fluorescence decay of each TRITC molecule was fitted using a double-exponential function. Throughout all the experiments, the intensity-weighted average fluorescence lifetime was obtained according to Equation (1):

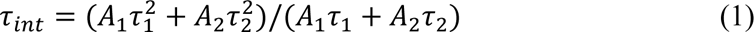

where *A* and *τ* represent the amplitude and fluorescence lifetime of each component, respectively. During observation, a liposome or aggregate of magnetic silica particles (Sicastar-M-CT, COOH, COREFRONT 6 μm; micromod Partikeltechnologie GmbH, Rostock, Germany) was illuminated using an 808 nm laser (530 or 950 μW, 10 s).

For cellular thermometry, iRGD-conjugated liposome containing calcein (10 μL) was dispersed into Dulbecco’s Modified Eagle’s Medium (DMEM) (190 μL). HeLa cells were incubated overnight in medium containing liposomes to facilitate the attachment of liposomes to the cell surface. Cells were then stained with FLIM-based thermometers, including PTG (plasma membrane), ETG (ER), and NTG (nucleus), at a final concentration of 500 nM (37 °C, 5 % CO_2_ for 30 min) (34). FLIM images of the PTG, ETG, and NTG were obtained using a 485 nm pulsed laser and a filter (520/35 Bright Line HC; Semrock). Data were analyzed using the aforementioned procedure. During observation, a single liposome was illuminated by an 808 nm laser (530 or 950 μW).

### Microscopic experiments for ACh release using VPc-Lipo encapsulating ACh

HEK293T cells expressing the GACh2.0 biosensor were prepared as previously described (45). Briefly, for transfection, HEK293T cells were incubated with the plasmid (pGACh2.0) and poly(ethylene imine) (PEI) in DMEM for 6 h, followed by washing with PBS and incubation in fresh DMEM for 48 h. The pDisplay-GACh2.0 was a gift from Yulong Li (Addgene plasmid 106073; Addgene, Watertown, MA, USA). To construct a calibration curve between the fluorescence intensity of the GACh2.0 biosensor and ACh concentration, fluorescence images were obtained at 256 × 256 pixels every 0.56 s using confocal microscopy (FV1200; Olympus Corporation). The same microscopic settings were used to study calcein release. During observation, DMEM containing ACh (at a final concentration of 0.1–1,000 μM) was sequentially added to the cells. To evaluate ACh release from the liposomes, HEK293T cells expressing GACh2.0 were incubated overnight in DMEM containing iRGD-conjugated ACh liposomes. The targeted liposomes attached to the cell membrane were illuminated with an 808 nm laser (950 μW, 50 ms) to release ACh. Fluorescence images were obtained according to the above-mentioned procedure.

### Modulation of Ca^2+^ dynamics in C2C12 myotubes and Drosophila brains

Myotubes differentiated from C2C12 myoblasts were incubated with DMEM containing VPc-Lipo-encapsulating ACh (10 μL) overnight. After washing with PBS, cells were stained with Fluo-4AM at a final concentration of 250 nM. Fluorescence images of Fluo-4 were captured at 256 × 256 pixels every 0.56 s. The microscopic settings were the same as those for the FITC channel (emission filter set: 490–540 nm; laser: 473 nm). During observation, the liposomes were illuminated by an 808 nm laser at a power of 950 μW for 50 ms several times to release ACh.

We followed previous studies for *ex vivo* application using the *Drosophila* brain, including imaging procedures (45). Briefly, flies (*Drosophila melanogaster*) were maintained on standard fly medium under normal conditions at 25 °C. Using the GAL4/UAS system, jGCaMP7c (BL#79030) as a calcium indicator was sparsely expressed in fruitless-expressing neurons (BL#66696). Eclosed male flies were maintained in groups (10–20 flies per vial). Throughout all experiments, flies were used 6–7 days after eclosion. All imaging experiments were performed using adult hemolymph-like (AHL) saline. Brain samples were dissected in Ca^2+^-free AHL saline, and the blood– brain barrier was digested with papain (10 U/mL) (Worthington Biochemical Corporation) for 15 min at room temperature. Brain samples were treated with AHL saline containing VPc-Lipo encapsulating ACh for 5 min. For the inhibition experiments, α-bungarotoxin (5 μM) or donepezil (100 nM) was added prior to imaging studies. Fluorescence images were acquired at 512 × 512 pixels every 1.64 s. The fluorescence of jGCaMP7c was accumulated in the FITC channel (emission filter set: 490–540 nm; laser: 473 nm). During observation, the liposomes were illuminated by an 808 nm laser at a power of 950 μW for 50 ms several times to release ACh.

## Author contributions

Conceptualization: SRS, TY, KS, IT, SA

Methodology: TA, IT, SH, TK, SRS, TY, YTC

Investigation: TY, SRS, IT, YK, KN, MI, TS, TF, AK

Visualization: TY, SA

Supervision: SA

Writing—original draft: TY

Writing—review & editing: all authors

## Data and materials availability

All data are available in the main text or the supplementary materials.

## Supplementary Materials

It is available via online.

