## Supplementary information for "Harnessing Dye-induced Photothermal Confinement in Lipid Membranes: A Path to NIR-modulated Artificial Synaptic Vesicles"

**Characterization of PTD embedded liposomes**

The photochemical parameters of PTDs were analyzed as follows. Time-resolved photoluminescence lifetime (τ) measurements were carried out using a time-correlated single photon counting lifetime spectroscopy system (Quantaurus-Tau C11367-22, Hamamatsu photonics). The decay constants and fitting parameters for the transient decays were determined using the software embedded in the Quantaurus-Tau system. Absolute fluorescence quantum yields (ΦPL) were measured using a calibrated integrating sphere system (Quantaurus-QY Plus C13534-22, Hamamatsu photonics). The diameter of liposomes was characterized by Zetasizer ZSP (Malvern). The differential scanning calorimetry was performed by diamond DSC (PerkinElmer). The Cryo-TEM image was taken by Terabase company (outsourcing).

**MD simulation**

**1-1 Simulation model:**

We performed molecular dynamics (MD) simulation of a lipid bilayer membrane consist of DPPC and SA with dye molecules (VPc, IR792, C6T1, and Calcein) using the LAMMPS package [1]. The CHARMM General Forcefield (CGenFF) [2] in conjunction with the TIP3P water model [3] was used. Long-range Coulomb interactions were handled by the particle mesh Ewald (PME) method. The atomic charges of all molecules were assigned using ab initio method via Gaussian16 with B3LYP functionals and 6-31G (d, p) basis set level [4]. A membrane sandwiched between water was used as done in previous studies [5-7]. The lipid bilayer was composed of 128 molecules (115 DPPC and 13 SA) with approximately 8400 water molecules in the initial simulation boxes (64.0 Å, 64.0 Å, and 120.0 Å). The initial placement of the molecules was performed using VMD [8].

**1-2 Stability of dye molecules on the lipid bilayer membrane:**

We evaluated the stabilization energies by performing MD simulations that accounted for the insertion of dye molecules into the lipid bilayer membrane, using various initial molecular configurations. Three models were employed for the molecular configurations: in water (IW), at the surface of the membrane (SM), and inside the membrane (IM). For the IW and SM initial structures, the dye molecule was positioned 20 Å and 2 Å from the membrane surface, respectively. The dye molecule in the initial structure of IM was inserted in the center of the membrane. The MD simulation applied a time step of 1.0 fs, an equilibration time of 3 ns, and a simulation time of 2 ns for stabilization energy analysis in the NPT ensemble at 300 K and 1 atm. The stabilization energy was calculated based on the difference of the averaged total energy between IW and SM or IW and IM.

**1-3 Heat transportation from dye molecule to membrane:**

To evaluate the heat transfer from light-absorbing dye molecules to lipid membranes, we used non-equilibrium MD simulations to investigate the time constants involved in the kinetic energy transport. We used the configurations at 300 K, for which the stabilization energy was assessed, without temperature or pressure control, i.e., in the NVE ensemble. Subsequently, non-equilibrium MD simulations were carried out with the Langevin dynamics thermostat for the dye molecule at 350 K to mimic the exothermic effect. Note that the membrane and water molecules were still in the NVE ensemble. The non-equilibrium MD simulations also applied a time step of 1.0 fs and a simulation time of 1 ns before heating and 20 ns under heating. The kinetic energy of each atom in the MD simulation was used to calculate the temperature increase $\Delta T_{\mathrm{MD}}(t)$ at time $t$, using the following equation:

$$\Delta T_{MD}(t) = \frac{2}{3Nk_B}\Delta E_{kin}(t),$$

$$\Delta E_{kin}(t) = \sum_{i}^{N}\left(E_{kin}^{i}(t) - \langle E_{kin}^{i}\rangle_{\mathrm{EQ}}\right).$$

$N$ is the number of atoms corresponding to the membrane, $k_B$ is the Boltzmann constant and index $i$ is atom index. $\Delta E_{kin}$ represents the sum of the kinetic energies increased from the equilibrium state of all atoms composing the membrane, $E_{kin}^{i}(t)$-$<E_{kin}^{i}>_{\mathrm{EQ}}$.

The time constant for temperature increase was introduced with the fitting function of $\Delta T_{MD}(t)$ to avoid the thermal fluctuation. Here, we suppose an exponential function $f(t)$ as the membrane temperature to reach equilibrium state under heating with time constant $\tau$.

$$f(t) = \Delta T_0\left(1 - e^{-\frac{t}{\tau}}\right),$$

$\Delta T_0$ is the temperature difference. Here $\Delta T_0$ was set at 50 K as the difference between 350 K and 300 K. $\tau$ was obtained by least-squares fitting $\Delta T_{\mathrm{MD}}(t)$ to $f$(t). Finally, the heat transportation property from dye molecule to membrane can be discussed by comparing $\tau$ with several dye molecules and their configuration.

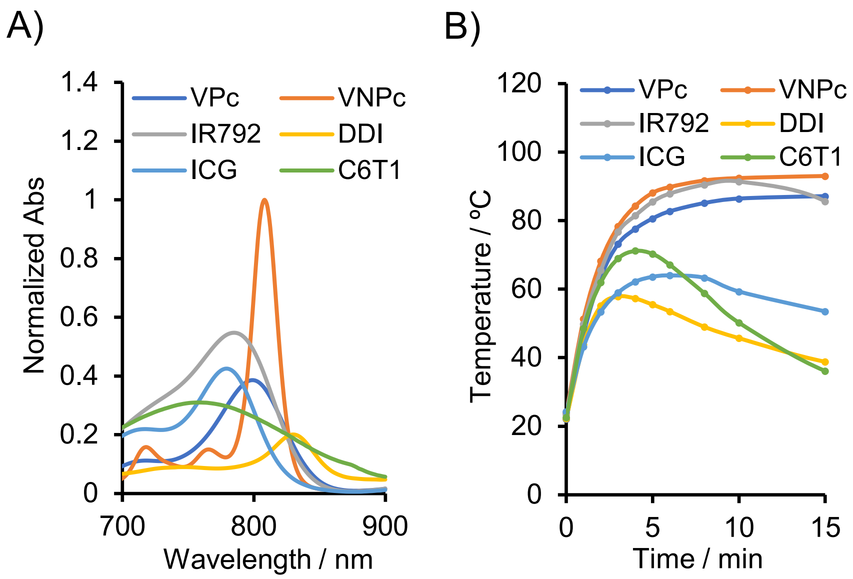


**Figure S1**. Optical properties of PTDs. A) Absorption spectra of PTDs in toluene. B) Heat production ability of PTDs in toluene under 808 nm laser illumination (160 mW).

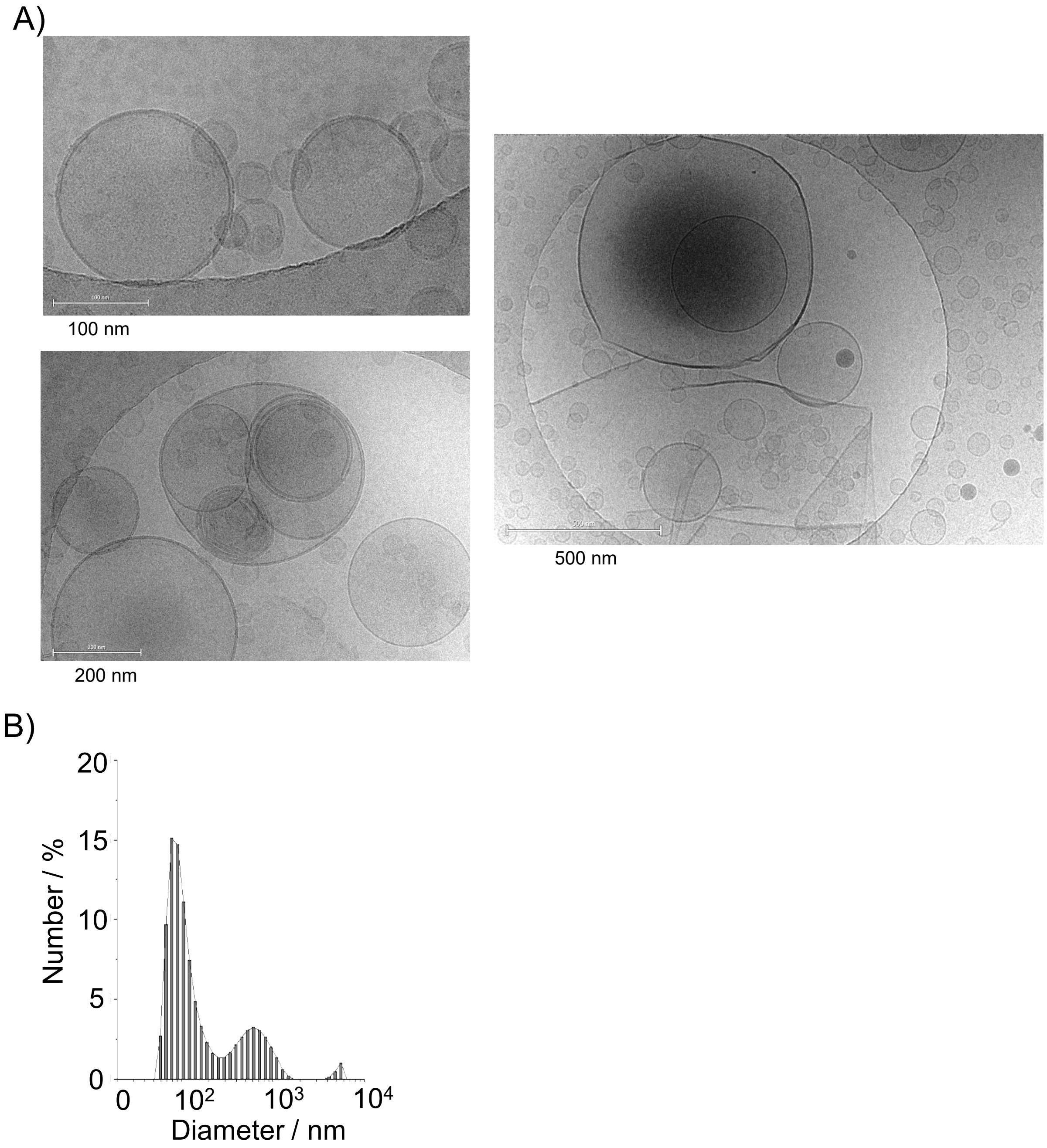


**Figure S2**. Characterization of the size of VPc embedded liposomes (VPc-Lipo). A) Transmission electron microscopy (TEM) revealing the uni-lamella structure of the liposomes, ranging from 50 nm to 500 nm in diameter. B) Distribution of particle diameter measured by dynamic light scattering (DLS).

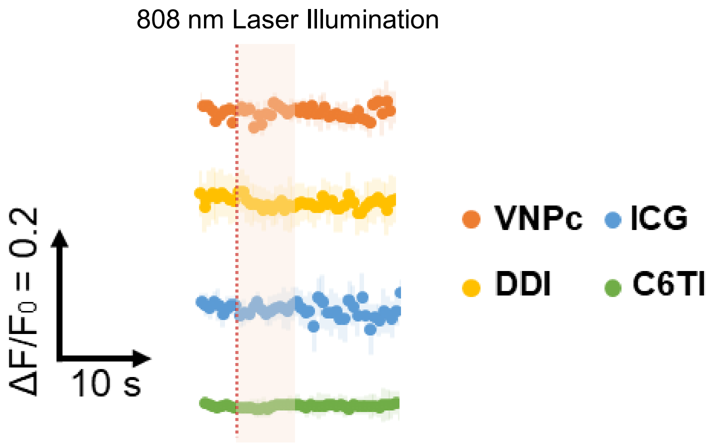


**Figure S3**. Investigation of PTD-embedded liposomes at microscopic level. The mean of calcein releases ($F/F_0$) with SD (n = 10) in VNPc, ICG, DDI and C6TI embedded liposomes were plotted in the time course. The red-dashed line represents the timing of 808 nm laser illumination (10 sec, 950 μW). The data is relevant to Fig. 2D.

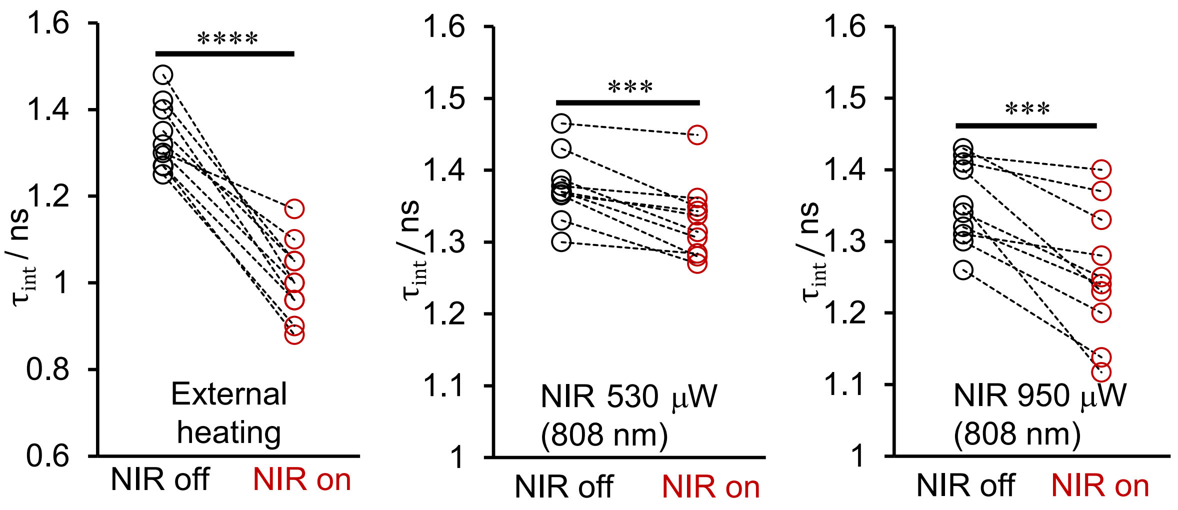


**Figure S4**. FLIM thermometry during local heating in the vicinity of the lipid bilayers of VPc-Lipo. FLIM thermometry was performed in the vicinity of the lipid bilayer of VPc-Lipo using tetramethylrhodamine (TRITC) as a FLIM temperature sensor, as shown in Fig. 4F. FLIM thermometry was conducted at the bilayer surface of VPc-Lipo under external heating (magnet particles) and local heating (808 nm laser: 530 and 950 μW) (n = 10 for each condition). Paired t-tests indicated significant differences (****p <0.0001, ***p <0.01). Based on these data, ΔT were estimated from the calibration curve presented in Fig. 4 (10 ± 2.4 ºC (external heating), 1.1 ± 1.4 ºC (530 μW, local heating), and 3.1 ± 2.5 ºC (950 μW, local heating).

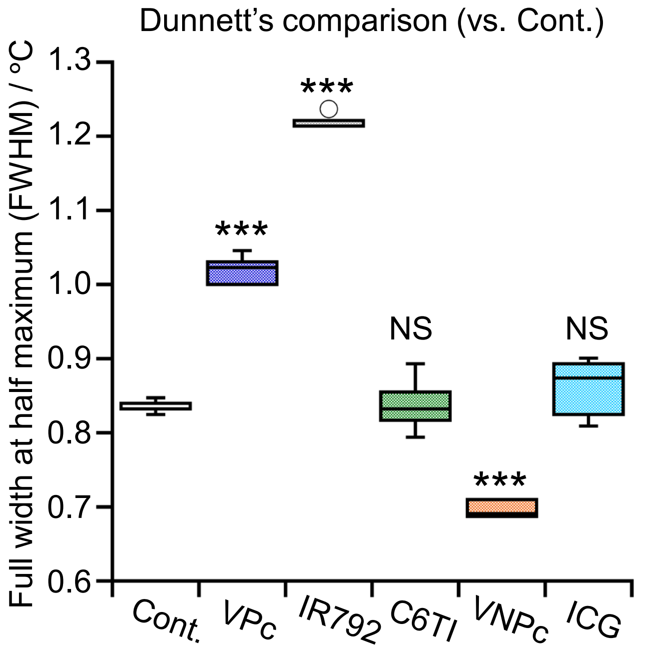


**Figure S5**. Full width at the half maximum measured by differential scanning calorimetry (DSC). The same lipid formulation was used as shown in Fig. 1. Each data point represents means ± SD (n = 6). Dunnett's multiple comparison test indicated significant differences (***p< 0.0001).

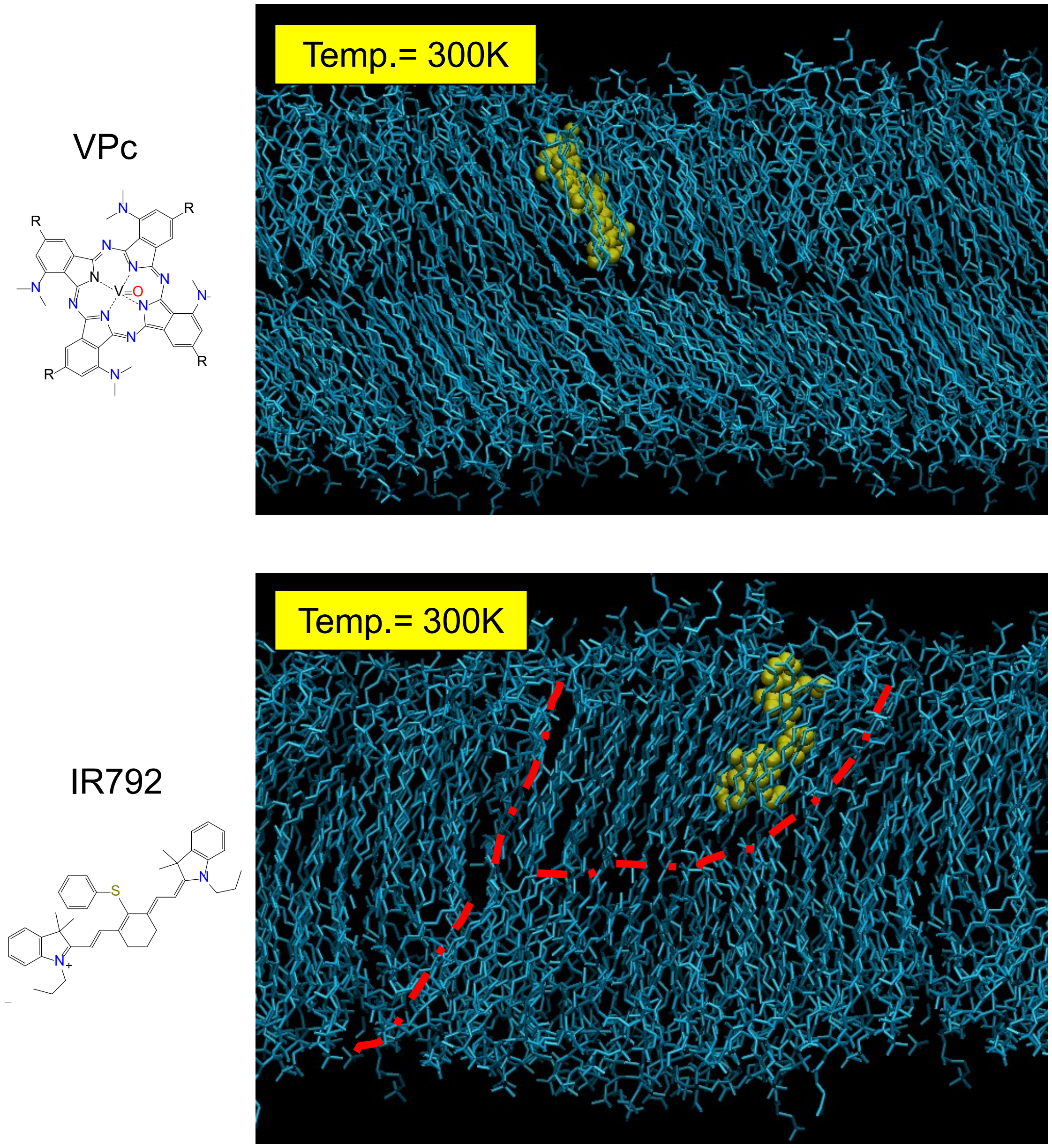


**Figure S6**. MD simulation of VPc and IR792 in a mixture of lipids. Following the insertion of the dye into the lipid membrane, VPc minimally disturbed the structure of lipid bilayers whereas IR792 generated cracking (red dashed line indicates the cracking). Such cracking could potentially interfere with effective heat dissipation.

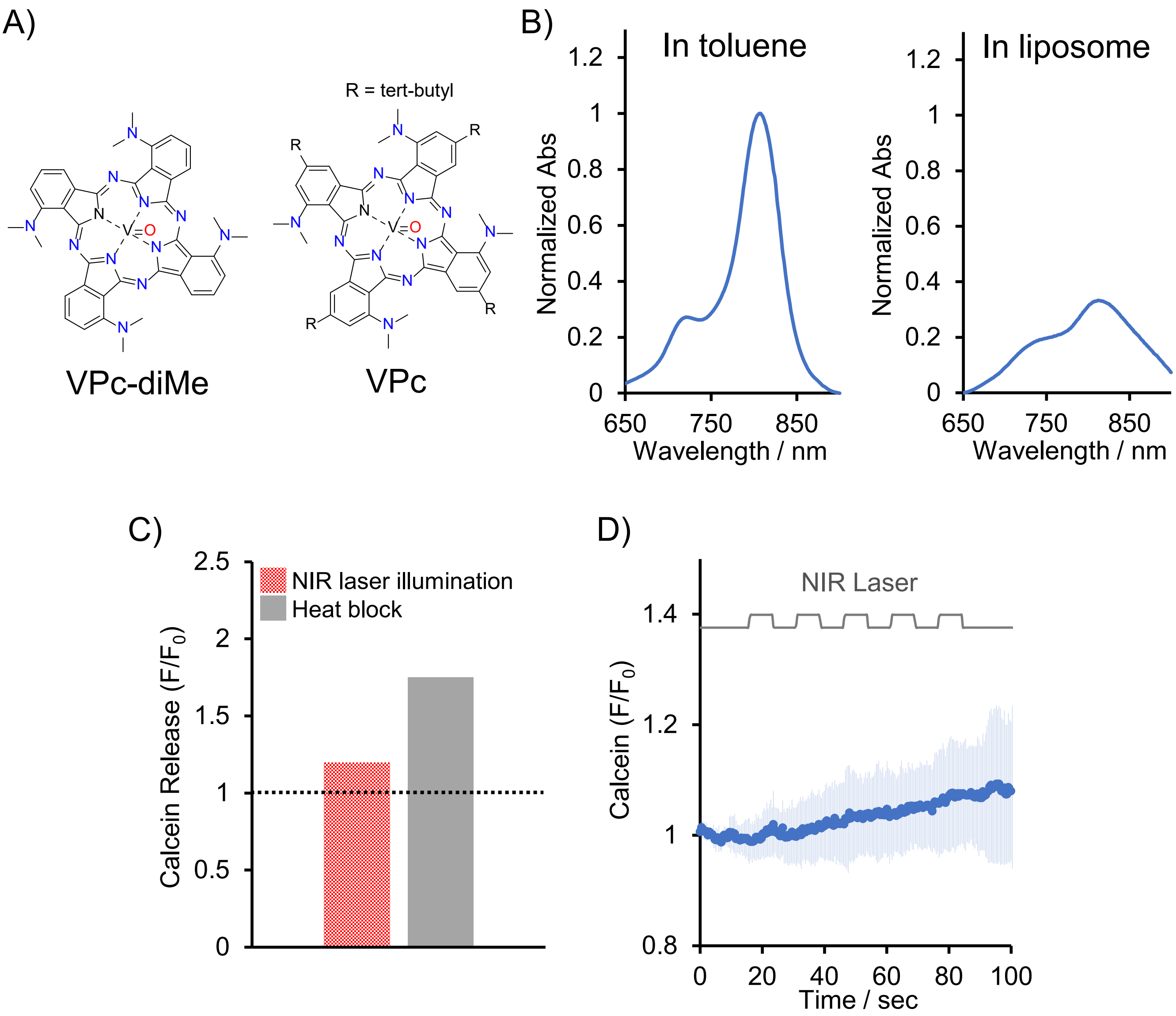


**Figure S7**. Structural relationship investigation of VPc-diMe without the tert-butyl group. A) Chemical structures of VPc-diMe (with VPc provided as a reference). B) Absorption spectra of VPc-diMe in toluene and liposome (DPPC/SA). C) Analysis of heat-induced leakage of calcein from VPc-diMe embedded liposomes. Liposome suspensions were heated with a heat blocker at 50 ºC for 20 min or 808 nm laser illumination (160 mW) for 5 min (same experiments with Fig. 1G). Changes in normalized fluorescence intensity of calcein ($F/F_0$) was evaluated in both heating methods. The dotted line represents a control without heating treatments ($F/F_0 = 1$ indicates no-leakage of calcein). D) Sequential 808 nm laser illumination (5 cycles, 10 sec, 950 μW) of VPc-diMe embedded liposome encapsulating calcein.

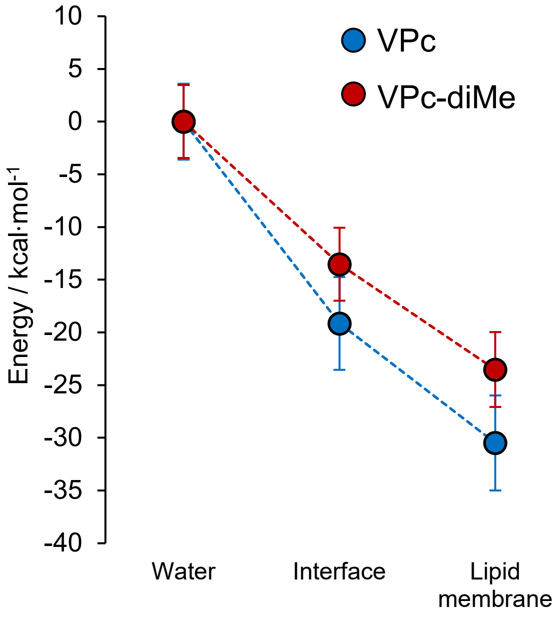


**Figure S8.** The potential energy of VPc and VPc-diMe was evaluated relative to their energy in water, with that data variation derived from each time step (0.5 ps) over 1 ns.

**Table S1**. Comparison with NIR-liposomal system and photocaged compounds

| | **Photo-caged compound** | **NIR-liposomal system** |
|---|---|---|
| Varieties of Compounds | Limited number available | A huge varieties of hydrophilic compounds |
| Spatial Resolution | Limited to laser spot size | Control by the liposome size |
| Temporal Resolution | From msec to sec | **msec** (independent of compounds) |
| Toxicity | Photo-toxicity (UV-vis range) | **Negligible** thermal damage |
| Solubility of compound | μM order | From **μM** to **mM** order |
